# PanTax: Strain-level taxonomic classification of metagenomic data using pangenome graphs

**DOI:** 10.1101/2024.11.15.623887

**Authors:** Wenhai Zhang, Yuansheng Liu, Jialu Xu, Enlian Chen, Alexander Schönhuth, Xiao Luo

## Abstract

Microbes are omnipresent, thriving in a range of habitats from oceans to soils and even within our gastrointestinal tracts. They play a vital role in maintaining ecological equilibrium and promoting the health of their hosts. Consequently, understanding the strain diversity within microbial communities is crucial, as variations between strains can lead to distinct phenotypic expressions or diverse biological functions. However, current methods for taxonomic classification from metagenomic sequencing data have several limitations, including their reliance solely on species resolution, support for either short or long reads, or their confinement to a given single species. Most notably, the majority of existing taxonomic classifiers rely solely on a single linear representative genome as a reference, which fails to capture the strain diversity, thereby introducing single-reference biases.

Here, we present PanTax, a pangenome graph-based taxonomic classification method that overcomes the shortcomings of single-reference genome-based approaches, because pangenome graphs possess the capability to depict the genetic variability present across multiple evolutionarily or environmentally related genomes. PanTax provides a comprehensive solution to taxonomic classification for strain resolution, compatibility with both short and long reads, and compatibility with single or multiple species. Extensive benchmarking results demonstrate that PanTax drastically outperforms state-of-the-art approaches, primarily evidenced by its significantly higher precision or recall (at both species and strain levels), while maintaining comparable or better performance in other aspects across various datasets. PanTax is a user-friendly open-source tool that is publicly accessible at https://github.com/LuoGroup2023/PanTax.

## Introduction

Microorganisms are ubiquitous on Earth, inhabiting diverse environments such as oceans, soils, and gastrointestinal tracts, where they play indispensable roles in maintaining ecological balance and host health. Microbial communities are often composed of multiple types of microorganisms, at uneven species abundance and high diversity. Rapid mutation and horizontal gene transfer in microorganisms may result in different strains of the same species, which further increases the biological diversity and complexity of microbial communities. Strain diversity of microbial communities is a key factor in microbiome-related research, as it does not only reflect the evolution and adaptation of microorganisms, but also their interactions with the environment and the functions of their host. Previous studies have shown that different strains may exhibit different phenotypes or perform different biological functions in the environment (Marx, 2016; Van Rossum *et al*., 2020). Moreover, recent works have revealed that strain-level genome variations in the gut microbiota are closely related to host health and disease (Zhernakova *et al*., 2024; Chen *et al*., 2022; Zeevi *et al*., 2019). This means that determining the composition of gut microbiota at the level of strains can accurately predict intestinal diseases such as inflammatory bowel disease (Jiang *et al*., 2022). This points out why deciphering the strain diversity of microbial communities is of great significance for understanding the biological characteristics of microorganisms in general and their involvement in scenarios of clinical interest in particular (Bickhart *et al*., 2022; Zheng *et al*., 2022; Fedarko *et al*., 2022). Therefore, to reveal the structure and function of microbial communities, we need to estimate their composition and abundance at high resolution, that is, not only at species level, but also at the level of strains.

Metagenome sequencing preserves the preserves the majority of the genetic information and enables a strain-level analysis of microbial communities. This establishes significant advantages over amplicon sequencing and culture-based methods, because these two approaches, by design, inevitably miss large parts of the genetic material one seeks to analyze. Next generation sequencing (NGS) has been routinely used in reference-based taxonomic classification of metagenomic data, which explains the development of many related tools. The fundamental tasks are to identify and classify all bacterial species and the corresponding strains that make part of a metagenome *(taxonomic binning)*, and to estimate the abundances of the participating species and their strains *(taxonomic profiling)*. Various tools have been developed that operate at the level of species alone. However, strain-level analysis still has remained a tough challenge. All of thes species-level tools receive the sequenced reads of a metagenome as input, and output the spectrum of species that make part of the metagenome (binning), and also possibly their relative abundances (profiling). Note that none of these tools aims to assemble the individual genomes prior to raising the relevant estimates. Rather, they put the reads of a metagenome into context with existing reference sequences, and derive the corresponding conclusions from the resulting read-to-reference sequence alignments.

Existing tools can be categorized into three types. The first are marker-based methods, such as MetaPhlAn2 (Truong *et al*., 2015), MetaPhlAn4 (Blanco-Míguez *et al*., 2023) and mOTUs2 (Milanese *et al*., 2019). These methods only use a subset of genes, usually particular gene families by which one can distinguish species. For instance, the 16S rRNA sequence is the most common single marker gene for bacterial metagenomics due to its high conservation (Edgar, 2018). While these methods are computationally efficient, the classification may be skewed if the marker genes in the microbial sequences of interest do not follow a uniform distribution (D’Amore *et al*., 2016). The second are DNA-to-protein methods, where DIAMOND (Buchfink *et al*., 2015), Kaiju (Menzel *et al*., 2016) and MMseqs2 (Steinegger and Söding, 2017) are prominent examples. DNA-to-protein methods focus on the protein-coding sequences of the genomes, while discarding all non-coding sequence from further consideration. This prevents the monitoring of differences that occur in the non-coding portions of the bacterial genomes. The third are DNA-to-DNA methods, such as Kraken (Wood and Salzberg, 2014), Kraken2 (Wood *et al*., 2019), KrakenUniq (Breitwieser *et al*., 2018), CLARK (Ounit *et al*., 2015), Centrifuge (Kim *et al*., 2016), Bracken (Lu *et al*., 2017) and CAMMiQ (Zhu *et al*., 2022), characterized by considering whole-genome information for classification. However, they usually select a single complete genome as representative for each species, which is unable to capture the genome diversity of various strains within a species. Therefore, these methods fail to classify reads that are highly divergent with the representative genomes, thus affecting the final classification performance. In addition, most of these methods only perform taxonomic classification at the species level, so do not operate at the level of strains. Strain-level microbiome composition analysis tools that have been suggested recently, such as StrainScan (Liao *et al*., 2023), StrainGE (van Dijk *et al*., 2022) or StrainEst (Albanese and Donati, 2017), can only handle one species specified by the user as input. They can only handle one species that preassigned by users. When considering these tools, it is important to understand that they particularly cater to the pecularities of metagenomes, which distinguishes them from tools that target the classification of isolate genomes, for example raised on Petri dishes or retrieved from environments that do not support entire microbial communities. See ProPhyle (B̌rinda *et al*., 2017) and Phylign (B̌rinda *et al*., 2023) for such approaches.

NGS or short-read sequencing, characterized by reads of a few hundred base pairs (bp) in length, has been the preferred method for metagenomics due to its ubiquitouos availability and its low cost. In the meantime, third generation sequencing (TGS) or long-read sequencing technologies such as Pacific Biosciences (PacBio) and Oxford Nanopore Technologies (ONT), have become more affordable. TGS has three advantages over short-read sequencing when dealing with metagenomic assignment and estimating the bacterial composition. First, long reads, of length tens of Kbp to Mbp, retain much longer-range genomic information. Therefore, they can more easily distinguish intra-genomic repeats or highly similar strains, which supports the disambiguation of reads when determining the strain they stem from. Second, as per the basic properties of single-molecule sequencing technologies, TGS (PacBio and ONT) avoid the generation of PCR duplicates common to NGS, which avoids the corresponding biases during taxonomic profiling. Third, portable TGS sequencers, such as ONT Flongle and MinION, enable cost-effective, in-field and real-time metagenomic sequencing. This can be used in scenarios where speed of identification of microbial/viral components matters, such as, for example, during pandemics (Quick *et al*., 2016; Brinkmann *et al*., 2021). Urgency may also be a factor in scenarios where samples are difficult to culture or preserve (Runtuwene *et al*., 2019; Wang *et al*., 2021), or when analyzing samples raised in hospitals (Chng *et al*., 2020). Still, the largest part of taxonomic classifiers is based on NGS. Since the corresponding built-in algorithms are not suitable for querying sequences that are long and/or affected by high error rates, their underlying technology cannot be straightforwardly adapted to processing TGS reads.

The difficulties involved in the adaptation of tools from NGS to TGS, as just outlined, explains why developing novel approaches for TGS-based taxonomic classification is imperative. So far, only few related tools have been designed, such as Melon (marker-based) (Chen *et al*., 2023), MEGAN-LR (DNA-to-protein) (Huson *et al*., 2018) and MetaMaps (DNA-to-DNA) (Dilthey *et al*., 2019). Just as for the tools already listed above, the characteristic drawbacks of the above-mentioned three categories apply also here. When raising a summarizing overall account of the characteristics of tools (see Table 1 for an overview), Centrifuger (Kim *et al*., 2016) emerges as the only method that can handle both NGS and TGS reads and perform strain-level taxonomic classification (binning and profiling) for multiple species at a time, so proves to be the only tool already available to address all currently driving issues in a universal manner. Still, however, Centrifuger is based on linear reference sequence when binning and profiling metagenomes, which incurs the disadvantages in terms of biases that we have mentioned above.

**Table 1.**
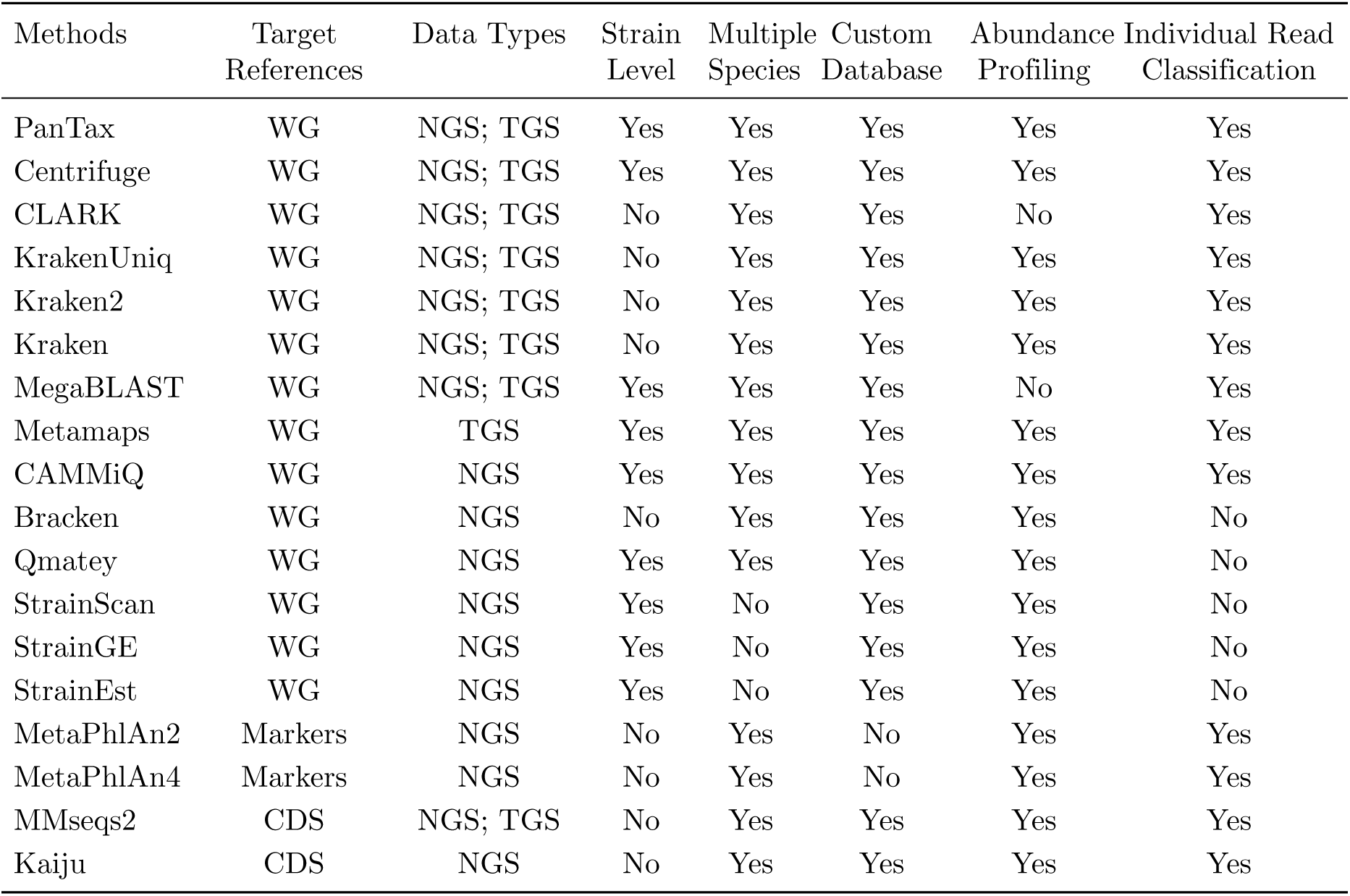
Characteristic summary of representative taxonomic classifiers. Note that Kraken, Kraken2, KrakenUniq, Bracken, Qmatey and MMseqs2 provide sequence abundances (i.e. how many reads are assigned to each taxon) instead of taxonomic relative abundances. WG: whole genome. CDS: coding sequence.

To overcome the shortcomings of existing taxonomic metagenome classifiers, we suggest PanTax ([Pan]genome graph based [Tax]onomic classifier), as a new method to perform strain-level taxonomic classification for both short-read and long-read metagenomic data. From a broader perspective, PanTax refers to the pangenome that captures the genomes of multiple strains as a reference, instead of only a single reference genome for each species. The corresponding expansion in terms of strain diversity that PanTax builds on enhances both the accuracy in mapping metagenome reads and the subsequent estimation of its composition at the level of strains. From a methodological point of view, PanTax’s innovation is the systematic integration of pangenome graphs, instead of plain linear sequences, as an algorithmic foundation for taxonomic classification. The particular type of pangenome graphs that PanTax relies on are variation graphs, as originally described in (Paten *et al*., 2017). In the meantime, variation graphs have proven effective in addressing a wide range of computational genomics challenges. Examples are primary read mapping and variant calling (Garrison *et al*., 2018; Martiniano *et al*., 2020; Siŕen *et al*., 2021), haplotype modeling (Rosen *et al*., 2017), as well as the refined correction of errors in long reads (Luo *et al*., 2022b). For the latter, greatest improvements were observed for long reads stemming from metagenomes. The favorable usage of variation graphs in the assembly of genomes from mixed samples (Baaijens *et al*., 2019, 2020) is an additional hint to the particular advantages of variation graphs in the analysis of metagenomes. To the best of our knowledge, PanTax is the first approach that employs pangenome graphs as reference systems for the taxonomic classification of metagenomes.

To provide evidence of PanTax’s superiority, we have conducted various challenging experiments and compared PanTax with the state-of-the-art taxonomic classifiers on both simulated and real datasets that relate to questions of actual interest in research. The corresponding benchmarking experiments consistently demonstrate that PanTax achieves the best performance rates in taxonomic classification, across all popular sequencing platforms.

## Results

We have developed and implemented PanTax, a novel method for classifying the contents of metagenomes. PanTax is universal insofar as it 1) evaluates whole genomes, 2) can take both NGS or TGS reads as input, 3) can process multiple species simultaneously, 4) refers to a custom database, which avoids the biases of external databases, 5) provides estimates on the abundances of the involved strains, 6) assigns single reads to strains, and, likely even most importantly, 7) operates at the level of strains. As was pointed out on plenty of occasions, all of these points are of critical relevance in the analysis of metagenomes. As can be seen from Table 1, Centrifuge (Kim *et al*., 2016) is the only tool that rivals PanTax in terms of covering this spectrum of essential qualities. We will demonstrate in the following, that PanTax outperforms Centrifuge across all relevant benchmarking criteria.

The foundation for PanTax’s superiority is to employ pangenome graphs, instead of linear genome sequences as a basis for the required reference systems. Unlike linear genome sequence, pangenome graphs, and here variation graphs in particular, have the capability to capture the full range of genetic variability characteristic of an evolutionarily or environmentally related collection of genomes (Paten *et al*., 2017). PanTax exploits this quality of variation graphs for capturing the genetic variation that distinguishes the different strains of a species in particular. In addition to this principled quality, pangenome graphs simply offer a considerably more compact way of representing multiple genomes in an encompassing context (Garrison *et al*., 2018; Siŕen *et al*., 2021). The practical advantages of pangenome graphs are the preservation of the strain-level diversity of metagenomes and the lack of ambiguity inducing redundancies by which read binning based approaches are affected.

In the following, we first present a comprehensive summary of the workflow of PanTax. Subsequently, we assess its performance at both the level of species and strains in comparison with all prominent state-of-the-art approaches, by experiments using both simulated and real data.

### Approach

Fig. 1 illustrates the workflow of PanTax. Here, we provide an overview of the workflow. For full details, see the Methods section.

**Figure 1.**
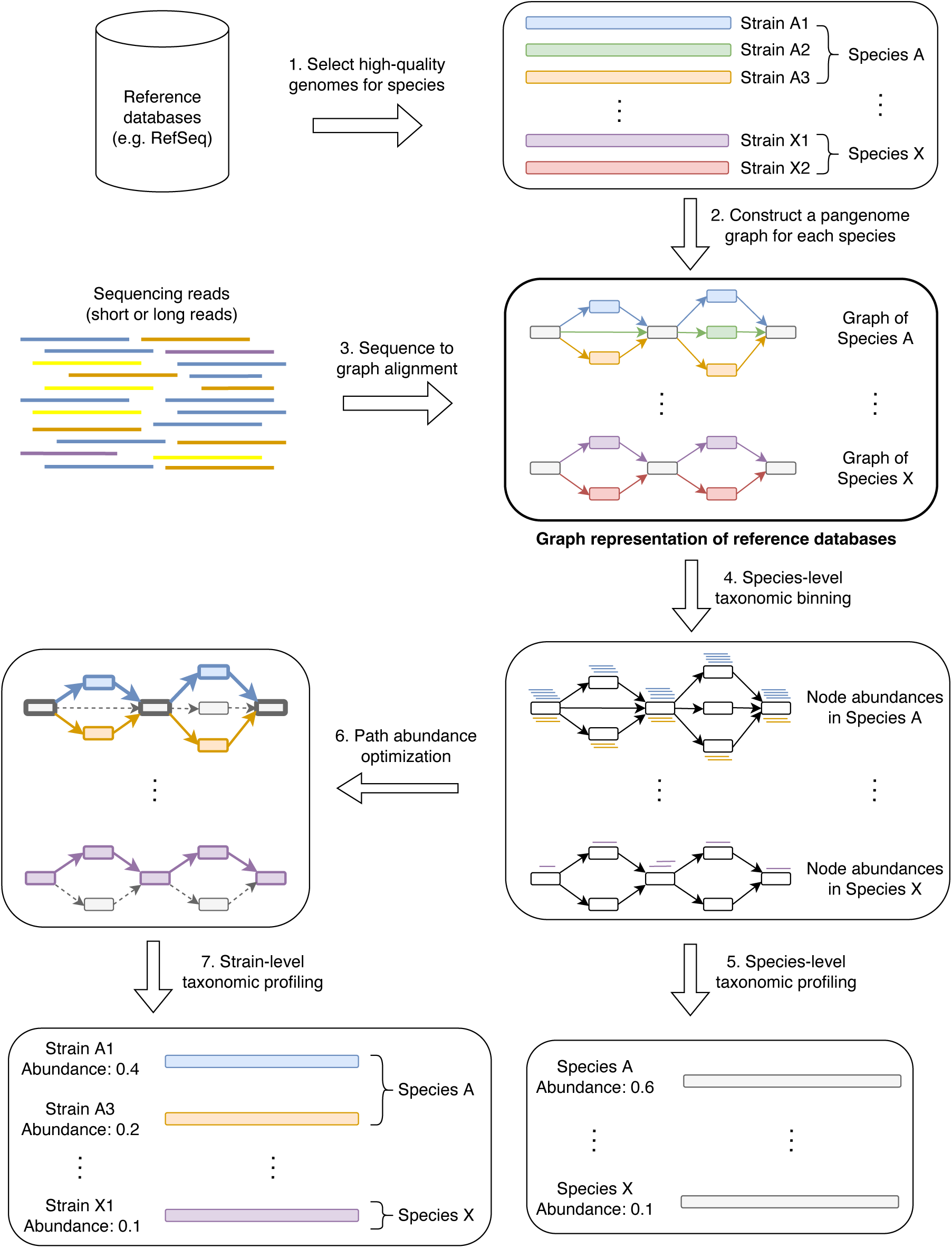
Workflow of PanTax. The color represents the strain specificity. The grey and colored rectangles in the pangenome graphs indicate the shared and unique genomic segments among strains, respectively. Arrows with the same color spell the path of a strain. The bold paths and dashed paths in the pangenome graphs indicate the existent and nonexistent strains in a sample, respectively. For simplicity, only two species (A and X) are shown in the figure. Note that step 6 and step 7 are optional unless one requires to perform strain-level taxonomic classification.

PanTax consists of three stages. The first stage is to construct a pangenome graph-based reference database (steps 1-2 in Fig. 1), where each species that makes part of the metagenome corresponds to a species-specific pangenome graph. The second stage (steps 3-5 in Fig. 1) yields species-level taxonomic classification results, by evaluating the amount of reads that gets aligned to each of the species graphs. The optional third stage eventually determines the strains and the corresponding relative abundances in a species (steps 6-7 in Fig. 1), based on the fact that the individual strains of a species correspond to the individual paths in a species pangenome graph.

From a technical point of view, the first stage reflects to construct a pangenome graph from a collection of high-quality strain-level genomes as stored in reference genome databases such as NCBI’s RefSeq(O’Leary *et al*., 2016), or, if available, in more specific, customized databases. The choice of genome collection can be determined by the user. The more refined collections there are available, the more accurate the results. Note that the flexibility in terms of database usage accounts for the fact that microbe genome databases are filling up rapidly; so, PanTax does not depend on a static snapshot of available microbial sequences, but is explicitly tailored to keep up with the rapid progress in this area of research.

Subsequently, for each read, one needs to determine the species-level graph that gives rise to the optimal read-to-graph alignment of that read. To considerably enhance this step, all species-level pangenome graphs are virtually merged into a large “pangenome super-graph” that captures all species that one has recorded in the first stage. The advantage of merging individual species graphs into one large graph is the fact that one has to align each read with this one large graph, instead of having to align it with each species graph. Apart from drastically decreasing the number of alignment operations, this also facilitates the evaluation of the possibly several alignments of an individual read with different species in a statistically unifying context.

PanTax supports alignment operations for both long and short reads. Based on the resulting alignments, the reads are classified at the level of species, a process referred to as *taxonomic binning* in linear genome based settings. We recall that the evaluation of the alignments can be performed in a consistent manner thanks to the fundamental quality of pangenome graphs to arrange related individual genomes in an encompassing context. Subsequently, the relative abundance of each of the species is estimated by evaluating the resulting (appropriately normalized) read coverages, which one refers to as *taxonomic profiling* in non-pangenome graph based settings. Eventually, if desired, strain-level classification can be performed by solving the path abundance optimization problem in each species-specific graph using linear programming (see Methods for definitions). Upon having estimated the abundance of each strain, potentially false positive strains are discarded.

### Datasets

#### Simulated datasets (sim-low, sim-high)

We made use of CAMISIM (Fritz *et al*., 2019) to generate 10 metagenomic datasets (2 × 5) with different complexities (i.e. low and high) and different sequencing read types (i.e. Illumina, PacBio HiFi/CLR and ONT R10.4/R9.4.1 reads), which reflect most of application scenarios in current metagenomic research. Here, we applied PBSIM2 (Ono *et al*., 2021), as a most recent tool to simulate PacBio HiFi and ONT reads instead of the default simulator (PBSIM) in CAMISIM using sampling-based simulation mode. The PacBio HiFi sample FASTQ file was down-sampled from an anaerobic digester sludge sequencing sample (SRA accession: ERR7015089) (Feng *et al*., 2022). The ONT R10.4 and R9.4.1 template files are derived from a recent ONT long-read metagenomic study (Sereika *et al*., 2022). The low complexity dataset (named “sim-low”) comprises 60 strains stemming from 30 species (so 2 strains per species on average), while the high complexity (named “sim-high”) dataset comprises 1000 strains from 373 species (so a little less than 3 strains per species on average). For all datasets, the genomes used were retrieved from RefSeq (O’Leary *et al*., 2016)(see Supplementary Data 1 for the details). For the sim-low dataset, the relative abundances of strains vary from 0.30% to 6.43% and corresponding sequencing coverage over different strains varies from 0.96x to 20.3x. The mean sequencing coverage of strains is about 5.3x. For the sim-high dataset, relative abundances of strains vary from 0.04% to 0.30%. The corresponding sequencing coverage over different strains vary from 1.0x to 8.8x, and the mean sequencing coverage of strains is about 2.9x. Notably, in real-world scenarios, some novel strains may be present in a sample, meaning that not all strains can be identified using the reference database. To simulate this situation, we have deliberately included some strains in our simulated samples that are not present in our pangenome reference databases. Specifically, for the sim-low dataset, 56 out of 60 strains are found in the pangenome reference databases, while the remaining 4 strains are not. Similarly, for the sim-high dataset, 795 out of 1000 strains are present in the pangenome reference databases, whereas the other 205 strains are absent.

#### Pre-simulated datasets (CAMI strain-madness, CAMI Gastrointestinal tract)

We also consider two pre-simulated samples of sequencing data from the 2nd CAMI Challenge (https://data.cami-challenge.org/cami2). Both datasets include short reads and long reads (PacBio CLR). For both, we selected the first sample (labeled with “sample 0”) from the respective community. The CAMI strain-madness dataset contains approximately 4 gigabytes of sequencing data, comprising 408 strains from 20 species, which reflects a high degree of richness in terms of strain diversity. The CAMI Gastrointestinal tract dataset also contains approximately 4G of sequencing data, but it comprises 69 strains from 26 species. Note that not all reads in the CAMI Gastrointestinal tract dataset can be assigned to the strain level, which is why we refrain from using it for strain-level benchmarking analysis.

#### Real datasets: ATCC, Zymo1, Zymo2, NWC

##### ATCC

The ATCC MSA-1003 mock community dataset, henceforth referred to as ATCC, encompasses sequencing data for both PacBio HiFi reads (SRA accession: SRR9328980, 39 gigabytes FASTQ) and Illumina reads (SRA accession: SRR8359173, 2.8 gigabytes FASTQ), which are publicly available on NCBI (Portik *et al*., 2022). This mock community is composed of 20 bacterial species with staggered abundances, evenly distributed across four tiers: 18%, 1.8%, 0.18%, and 0.02%, with five species per tier.

##### Zymo1

The ZymoBIOMICS Microbial Community Standard dataset (catalog number: D6330), referred to as Zymo1, comprises sequencing data for Oxford Nanopore Technologies (ONT) R10.4 reads (https://github.com/LomanLab/mockcommunity, 24 gigabytes FASTQ), ONT R9.4.1 reads (SRA accession: ERR3152364, 27 gigabytes FASTQ), and Illumina reads (SRA accession: ERR2984773, 6.2 gigabytes FASTQ) from an even mock community (Somerville *et al*., 2019). Note that we used both ONT R10.4 and R9.4.1 data from Oxford Nanopore GridION sequencing platform. This mock community comprises 10 microbial species: 8 bacteria and 2 yeasts. The relative abundance of each bacteria is 12%, while the relative abundance of each yeast is 2%. As we mainly focus on bacteria in this study, we omitted the two yeasts in our analysis. Therefore, the actual relative abundance of each of the 8 bacteria is re-normalized and designated as 12.5%.

##### Zymo2

The ZymoBIOMICS gut microbiome standard dataset (catalog number: D6331), henceforth referred to as Zymo2, encompasses sequencing data for PacBio HiFi reads (SRA accession: SRR13128014, 34 gigabytes FASTQ). This standard dataset simulates the human gut microbiome by comprising 17 species, including 14 bacteria, 1 archaea, and 2 yeasts, with staggered abundances. According to this study (Portik *et al*., 2022), five strains of *E. coli* (each at 2.8% abundance) are treated as a single species with a combined abundance of 14%. Due to species synonymy, *Lactobacillus fermentum* is considered as *Limosilactobacillus fermentum*. The abundance distribution is staggered, with five species at 14%, four species at 6%, four species at 1.5%, and one species each at 0.1%, 0.01%, 0.001%, and 0.0001%. Similarly to Zymo1, in Zymo2, only the 14 bacteria were utilized for the analysis, and their abundances were re-normalized to accurately reflect their true abundances.

##### NWC

Another real dataset, referred to as NWC herein, is a publicly available metagenomic dataset derived from natural whey starter cultures. As reported in the original study(Somerville *et al*., 2019), two NWC samples were sequenced, and we chose the sequencing data of the NWC 2 sample for our experiments. This dataset comprises PacBio reads (SRA accession: SRR7585901, 9.4 gigabytes FASTQ) and ONT reads (SRA accession: SRR7585900, 2.7 gigabytes FASTQ). The PacBio sequencing data was obtained using the Sequel platform (v.2.1), while the ONT sequencing data was sequenced on a FLO-MIN107 (R9.5) flow cell. While the true abundance of species in this dataset is unknown, the authors determined the relative abundance composition of the species by oligotyping the 16S rRNA amplicon data of the sample. This relative abundance distribution can serve as a proxy for true abundance and will be used for comparison with other methods. Specifically, *S. thermophilus* occurs at 45.4% abundance, *L. delbrueckii* at 26.1%, *L. helveticus* at 25.6%, and other unknown species account for 2.9%. Note that we only evaluated three known species in benchmarking experiments.

### Remarks on evaluating alternative methods

To ensure a rigorous comparison, we have comprehensively selected a variety of representative taxonomic binning and profiling methods (see Table 1) and evaluated them concurrently. Given the principled differences between NGS and TGS reads, we conducted comparisons separately for these two types of reads. Specifically, for species-level classification of multiple species, we ran representative tools such as CLARK, KrakenUniq, Kraken, Kraken2, Centrifuge, Bracken, MetaPhlAn4 and Qmatey on NGS sequencing datasets, and ran representative tools such as CLARK, KrakenUniq, Kraken, Kraken2, Centrifuge, MetaMaps and MegaBLAST on TGS sequencing datasets, respectively. For strain-level classification of multiple species, we ran Centrifuge and Qmatey on NGS sequencing datasets, while we ran Centrifuge and MetaMaps on the TGS sequencing datasets. Note that we excluded CAMMiQ from our benchmarking evaluations because runs ended in segmentation faults in all our attempts. As for strain-level classification of single species, we compared with the state-of-the-art methods StrainScan, StrainGE and StrainEst, all of which are specifically designed for this task.

Note that it was observed that usage of default reference databases can introduce systematic biases when using taxonomic classifiers (Simon *et al*., 2019). To mitigate this effect, we utilized a unified set of reference genomes for all methods involved, excluding the marker-based tools. When selecting species-specific reference genomes, we opted for the longest representative as part of the respective pangenome of the species. As selecting the longest possible genome maximizes the amount of sequence for each species, this decreases the odds to fail to align species-compatible sequence to a maximum degree.

We deal with two distinct reference databases for taxonomic classification. The first is called RefDB:13404, because it encompasses 13,404 strains(8778 species). We use RefDB:13404 for constructing pangenomes, because it lists several strains for many species. The second database is called RefDB:8778, because it comprises 8 778 strains. Unlike for RefDB:13404, each of the strains in RefDB:8778 is from a different species, so is the designated representative of its species. For species-level classification in the multiple species experiments, PanTax utilizes RefDB:13404. All other tools (with the exception of the marker-based tools), however, need to employ RefDB:8778 because they crucially rely on a single representative strain as a reference. For strain-level classification in the multiple species experiments, all methods refer to RefDB:13404, because one needs to include all strains into the reference databases for strain-level classification. For other scenarios that the reference database undergo reconstruction, additional instructions will be furnished accordingly.

For the obvious (fairness guided) reasons, we do not report results on the classification of individual reads for methods that solely support the estimation of relative abundances. Conversely, we do report results on relative abundances for methods that support the classification of individual reads, by estimating species abundance values via base counts derived from individual read counts the obvious way.

### Performance evaluation

To evaluate taxonomic classifiers, we rely on metrics like precision, recall and F1 score, which assess the performance in terms of the classification of reads and the identification of taxa. It is important to realize that these metrics tend to neglect low-abundance taxa due to these metrics may overlook low-abundance taxa. For a more comprehensive evaluation, we utilize precision-recall curves and the area under the curve (AUPR), which capture the classifier’s performance across all taxa, including those of low abundance. In addition, accurate estimation of taxon abundances in metagenomes is essential. We use L2 distance to quantify the abundance profile similarity. However, L2 distance is sensitive to high-abundance taxa. Therefore, we also introduce the absolute frequency error (AFE) and relative frequency error (RFE) as complementary metrics to capture abundance estimation performance more comprehensively. See “Metrics for evaluation” subsection in Methods for full details.

### Tasks

For taxonomic classification (binning and profiling) of metagenomic data, we focus on five computational tasks in this study, namely, species-level taxonomic binning, species-level taxonomic profiling, strain-level taxonomic binning (multiple species), strain-level taxonomic profiling (multiple species), and strain-level taxonomic classification(binning and profiling) for single species. Species-level taxonomic binning assigns reads to species, aiding in identifying microbial composition, while species-level profiling estimates their relative abundances. Strain-level taxonomic binning and profiling for multiple species provides a detailed view of strain diversity and abundance across different species, enhancing our understanding of microbial dynamics. Additionally, strain-level classification for a single species allows for in-depth analysis of strain variation within that species. We evaluate PanTax and other state-of-the-art methods using both NGS and TGS data from various datasets to ensure comprehensive benchmarking. See below for the details.

### Species level taxonomic binning

We recall that taxonomic classification involves two processes: taxonomic binning, that is the assignment of reads to taxa, and taxonomic profiling, which is the estimation of the (relative) abundances of the taxa present in the metagenome. In this subsection, we focus on species-level taxonomic binning. In other words, we evaluate methods in terms of their performance with respect to assigning individual reads to species. We evaluate all applicable state-of-the-art approaches on both NGS and TGS data from all datasets described above, which refers to pre-simulated, freshly simulated and real datasets.

#### Simulated datasets

As shown in Table 2, the benchmarking results for species-level taxonomic binning across two simulated datasets demonstrate that PanTax consistently achieves superior precision, recall, F1 score, and AUPR across all sequencing data types, including NGS, PacBio HiFi/CLR, and ONT R9.4.1/R10.4, with one exception: on NGS, PanTax only achieves second-best precision.

**Table 2.**
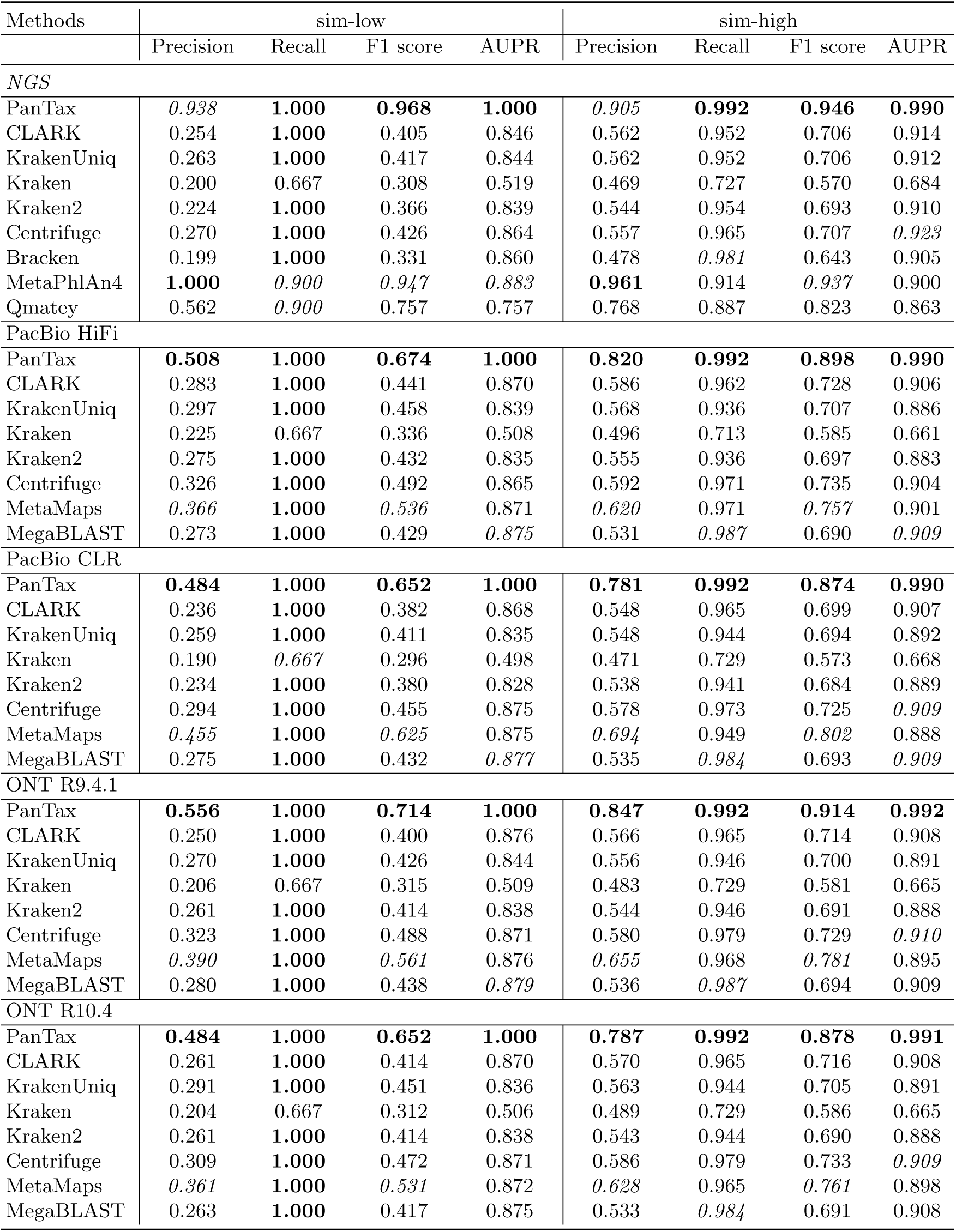
Benchmarking results of species-level taxonomic binning on the freshly simulated datasets (sim-low and sim-high). AUPR: area under the precision-recall curve. Note that the best score is marked in bold, and the second best score is marked in italics.

Specifically, for the sim-low dataset, PanTax attains precision ranging from 0.484 to 0.556 on TGS data, whereas other methods achieve only 0.190 to 0.455 (with MetaMaps having the second-best precision). Although PanTax secures the second-highest precision (0.938) on NGS data, trailing MetaPhlAn4 (precision=1.0), it surpasses MetaPhlAn4 in recall, F1 score, and AUPR. Additionally, PanTax achieves an AUPR of 1.0 on both NGS and TGS data, whereas other methods only reach an AUPR of 0.519 to 0.883 on NGS and 0.498 to 0.879 on TGS data.

A similar trend is observed in the sim-high dataset. Specifically, PanTax attains precision between 0.781 and 0.847 on TGS data, whereas other methods only manage 0.471 to 0.694 (with MetaMaps again achieving the second-best precision). While PanTax secures the second-highest precision (0.905) on NGS data, behind MetaPhlAn4 (precision=0.961), it outperforms MetaPhlAn4 in terms of recall, F1 score, and AUPR. Furthermore, PanTax achieves a recall of 0.992 on both NGS and TGS data, whereas other methods attain a recall of 0.727 to 0.981 on NGS and 0.713 to 0.987 on TGS data. Moreover, PanTax achieves an AUPR of approximately 0.99 on both NGS and TGS data, while other methods only achieve an AUPR of 0.684 to 0.923 on NGS and 0.661 to 0.910 on TGS data.

Notably, Supplementary Table 1 indicates that PanTax consistently achieves the highest precision for individual read classification across all simulated datasets; MetaPhlAn4 lacks this capability because it does not provide read-level classification results. Interestingly, Table 2 also shows that PanTax achieves better precision on NGS data compared to TGS data, while other methods exhibit similar precision on both types of data. Although this may seem counterintuitive, it can possibly be explained by the use of different sequence-to-graph algorithms for the two data types. Notably, PanTax achieves significantly better precision on higher complexity TGS data without compromising other metrics such as recall and AUPR.

#### Pre-simulated datasets

Table 3 presents the benchmarking results for species-level taxonomic binning across two pre-simulated datasets downloaded from the CAMI challenge.

**Table 3.** Benchmarking results of species-level taxonomic binning on the pre-simulated datasets (CAMI strain-madness and CAMI Gastrointestinal tract). AUPR: area under the precision-recall curve. Note that the best score is marked in bold, and the second best score is marked in italics.

| Methods | CAMI strain-madness |  |  |  | CAMI Gastrointestinal tract |  |  |  |
| --- | --- | --- | --- | --- | --- | --- | --- | --- |
|  | Precision | Recall | F1 score | AUPR | Precision | Recall | F1 score | AUPR |
| *NGS* |  |  |  |  |  |  |  |  |
| PanTax | 0.135 | **0.750** | 0.229 | *0.335* | *0.591* | **1.000** | *0.743* | **0.794** |
| CLARK | 0.049 | **0.750** | 0.092 | 0.234 | 0.325 | **1.000** | 0.491 | *0.732* |
| KrakenUniq | 0.062 | **0.750** | 0.115 | 0.254 | 0.366 | **1.000** | 0.536 | *0.732* |
| Kraken | 0.033 | 0.500 | 0.062 | 0.100 | 0.295 | 0.692 | 0.414 | 0.548 |
| Kraken2 | 0.051 | **0.750** | 0.096 | 0.244 | 0.342 | **1.000** | 0.510 | 0.718 |
| Centrifuge | 0.082 | **0.750** | 0.147 | 0.306 | 0.361 | **1.000** | 0.531 | 0.728 |
| Bracken | 0.054 | **0.750** | 0.101 | 0.206 | 0.361 | **1.000** | 0.531 | 0.710 |
| MetaPhlAn4 | **0.667** | 0.500 | **0.571** | **0.451** | **0.750** | 0.923 | **0.828** | 0.598 |
| Qmatey | *0.169* | *0.600* | *0.264* | 0.240 | 0.214 | 0.923 | 0.348 | 0.664 |
| *PacBio CLR* |  |  |  |  |  |  |  |  |
| PanTax | **0.112** | **0.750** | **0.195** | **0.353** | *0.426* | **1.000** | *0.598* | 0.739 |
| CLARK | 0.055 | *0.700* | 0.101 | 0.259 | 0.342 | **1.000** | 0.510 | *0.755* |
| KrakenUniq | 0.057 | *0.700* | 0.105 | 0.260 | 0.356 | **1.000** | 0.525 | 0.753 |
| Kraken | 0.036 | 0.600 | 0.068 | 0.101 | 0.281 | 0.692 | 0.400 | 0.567 |
| Kraken2 | 0.041 | *0.700* | 0.077 | 0.254 | 0.100 | **1.000** | 0.182 | 0.741 |
| Centrifuge | 0.031 | *0.700* | 0.060 | *0.292* | 0.064 | **1.000** | 0.120 | 0.692 |
| MetaMaps | *0.103* | 0.550 | *0.173* | 0.194 | **0.619** | **1.000** | **0.765** | **0.799** |
| MegaBLAST | 0.073 | 0.650 | 0.132 | 0.236 | 0.394 | **1.000** | 0.565 | 0.730 |

For the CAMI strain-madness dataset, PanTax achieves the third-highest precision (0.135) on NGS data, trailing MetaPhlAn4 (precision=0.667) and Qmatey (precision=0.169). However, PanTax significantly surpasses MetaPhlAn4 and Qmatey in terms of recall. On PacBio CLR data, PanTax demonstrates superior performance across all metrics, including precision, recall, F1 score, and AUPR. Supplementary Table 2 highlights that PanTax consistently achieves the highest precision and for individual read classification across both data types in the CAMI strain-madness dataset, a capability that MetaPhlAn4 and Qmatey lack since they do not provide read-level classification results.

In the CAMI Gastrointestinal tract dataset, PanTax attains the second-highest precision (0.591) on NGS data, lower than MetaPhlAn4 (precision=0.750), but it greatly surpasses MetaPhlAn4 in recall and AUPR. For PacBio CLR data, MetaMaps achieves the best performance across all metrics, while PanTax achieves the highest recall (1.0) and the second-best precision and F1 score. Notably, in read-level classification, PanTax achieves comparable precision (0.493 vs. 0.500) compared to MetaMaps, as shown in Supplementary Table 2.

#### Real datasets

Table 4 outlines the benchmarking outcomes for species-level taxonomic binning across four real datasets.

**Table 4.** Benchmarking results of species-level taxonomic binning on the real datasets (Zymo1, Zymo2, ATCC, NWC). Considering the presence of ultra-low abundance species, the threshold abundance of species reported on the Zymo2 HiFi dataset was set to 0 instead of 0.0001. MegaBLAST was killed after running for more than fifteen days. AUPR: area under the precision-recall curve. Note that the best score is marked in bold, and the second best score is marked in italics. N/A: not applicable.

| Methods | Precision | Recall | F1 score | AUPR | Precision | Recall | F1 score | AUPR |
| --- | --- | --- | --- | --- | --- | --- | --- | --- |
|  | Zymo1 NGS |  |  |  | Zymo1 ONT R9.4.1 |  |  |  |
| PanTax | 0.120 | 0.750 | 0.207 | *0.680* | 0.073 | **1.000** | 0.136 | **0.860** |
| CLARK | 0.077 | *0.875* | 0.141 | 0.558 | 0.038 | 0.875 | 0.073 | 0.513 |
| KrakenUniq | 0.080 | *0.875* | 0.147 | 0.549 | 0.067 | 0.875 | 0.124 | 0.450 |
| Kraken | 0.089 | *0.875* | 0.161 | 0.555 | 0.072 | 0.875 | 0.133 | 0.453 |
| Kraken2 | 0.060 | *0.875* | 0.113 | 0.544 | 0.053 | 0.875 | 0.101 | 0.441 |
| Centrifuge | 0.081 | **1.000** | 0.150 | 0.618 | 0.068 | **1.000** | 0.128 | *0.566* |
| Bracken | 0.070 | **1.000** | 0.131 | 0.429 | N/A | N/A | N/A | N/A |
| MetaPhlAn4 | **0.636** | *0.875* | **0.737** | **0.875** | N/A | N/A | N/A | N/A |
| Qmatey | *0.357* | 0.625 | *0.455* | 0.495 | N/A | N/A | N/A | N/A |
| MetaMaps | N/A | N/A | N/A | N/A | **0.129** | **1.000** | **0.229** | 0.353 |
| MegaBLAST | N/A | N/A | N/A | N/A | *0.082* | **1.000** | *0.151* | 0.460 |
|  | Zymo2 HiFi |  |  |  | Zymo1 ONT R10.4 |  |  |  |
| PanTax | **0.188** | **0.857** | **0.308** | **0.586** | 0.084 | **1.000** | 0.155 | **0.840** |
| CLARK | 0.095 | *0.786* | 0.169 | *0.579* | 0.039 | 0.875 | 0.074 | 0.423 |
| KrakenUniq | 0.115 | *0.786* | 0.200 | 0.531 | 0.064 | 0.875 | 0.119 | 0.372 |
| Kraken | 0.111 | 0.714 | 0.192 | 0.376 | 0.071 | 0.875 | 0.131 | 0.383 |
| Kraken2 | 0.116 | *0.786* | 0.202 | 0.519 | 0.053 | 0.875 | 0.101 | 0.364 |
| Centrifuge | 0.103 | *0.786* | 0.182 | 0.547 | 0.073 | **1.000** | 0.137 | 0.536 |
| MetaMaps | *0.131* | *0.786* | *0.224* | 0.532 | **0.130** | 0.875 | **0.226** | 0.355 |
| MegaBLAST | - | - | - | - | *0.092* | **1.000** | *0.168* | 0.370 |
|  | ATCC NGS |  |  |  | ATCC HiFi |  |  |  |
| PanTax | 0.295 | **0.900** | *0.444* | *0.568* | **0.198** | **0.800** | **0.317** | **0.540** |
| CLARK | 0.143 | 0.800 | 0.242 | 0.470 | 0.103 | *0.700* | 0.179 | 0.395 |
| KrakenUniq | 0.143 | 0.800 | 0.242 | 0.467 | 0.117 | 0.600 | 0.195 | 0.393 |
| Kraken | 0.127 | 0.650 | 0.213 | 0.348 | 0.100 | 0.550 | 0.169 | 0.280 |
| Kraken2 | 0.124 | 0.850 | 0.217 | 0.460 | 0.113 | 0.600 | 0.190 | 0.380 |
| Centrifuge | 0.145 | 0.850 | 0.248 | 0.483 | 0.121 | 0.650 | 0.205 | *0.399* |
| Bracken | 0.115 | 0.850 | 0.202 | 0.419 | N/A | N/A | N/A | N/A |
| MetaPhlAn4 | **0.875** | 0.700 | **0.778** | **0.666** | N/A | N/A | N/A | N/A |
| Qmatey | *0.714* | 0.250 | 0.370 | 0.250 | N/A | N/A | N/A | N/A |
| MetaMaps | N/A | N/A | N/A | N/A | *0.197* | 0.650 | *0.302* | 0.369 |
| MegaBLAST | N/A | N/A | N/A | N/A | - | - | - | - |
|  | NWC PacBio |  |  |  | NWC ONT |  |  |  |
| PanTax | 0.091 | **1.000** | 0.167 | **1.000** | **0.100** | **1.000** | **0.182** | **1.000** |
| CLARK | 0.067 | **1.000** | 0.125 | **1.000** | 0.079 | **1.000** | 0.146 | **1.000** |
| KrakenUniq | 0.077 | **1.000** | 0.143 | **1.000** | 0.073 | **1.000** | 0.136 | **1.000** |
| Kraken | 0.054 | 0.667 | 0.100 | 0.667 | 0.053 | 0.667 | 0.098 | 0.667 |
| Kraken2 | *0.100* | **1.000** | *0.182* | **1.000** | 0.079 | **1.000** | 0.146 | **1.000** |
| Centrifuge | 0.060 | **1.000** | 0.113 | **1.000** | 0.053 | **1.000** | 0.100 | **1.000** |
| MetaMaps | **0.130** | **1.000** | **0.231** | **1.000** | *0.094* | **1.000** | *0.171* | **1.000** |
| MegaBLAST | 0.083 | **1.000** | 0.154 | **1.000** | 0.060 | **1.000** | 0.113 | **1.000** |

On the Zymo1 dataset (NGS), MetaPhlAn4 excels in precision, F1 score, and AUPR but has lower recall. PanTax, on the other hand, achieves the third-highest precision and F1 score, along with the second-best AUPR. For the Zymo1 dataset (ONT R9.4.1/R10.4), PanTax demonstrates the highest recall (1.0/1.0) and AUPR (0.86/0.84), and the third-highest precision (0.073/0.084). Although its precision is lower than MetaMaps (0.129/0.130), PanTax significantly outperforms MetaMaps in recall (1.0/0.875) and AUPR (notably 0.353/0.355).

On the Zymo2 dataset (PacBio HiFi), PanTax achieves the best performance in all metrics, including precision, recall, F1 score, and AUPR.

For the ATCC dataset (NGS), PanTax ranks third in precision (0.295), following MetaPhlAn4 (precision=0.875) and Qmatey (precision=0.714). Nevertheless, PanTax surpasses MetaPhlAn4 and Qmatey in recall. On PacBio HiFi data, PanTax outperforms all other methods across all metrics, including precision, recall, F1 score, and AUPR.

In the NWC dataset (PacBio), PanTax secures the third-highest precision (0.091), behind MetaMaps (precision=0.13) and Kraken2 (precision=0.10), with similar recall and AUPR scores. On ONT data, PanTax achieves the highest precision (0.1), outperforming MetaMaps (precision=0.094) and all others, while all methods are on a par with each other for the other metrics Recall and AUPR, where everyone achieves (near-)optimal scores.

### Species level taxonomic profiling

The taxonomic profiling process aims to compute the relative abundances of taxa (i.e. species or strains) present in metagenomic samples. We also evaluated those representative tools using both NGS and TGS data on various simulated, readily available pre-simulated, and real datasets.

#### Simulated datasets

As illustrated in Table 5, the benchmarking results for species-level taxonomic profiling across two simulated datasets reveal that PanTax consistently achieves the lowest L2 distance across all sequencing data types, including NGS, PacBio HiFi/CLR, and ONT R9.4.1/R10.4. For instance, PanTax achieves an L2 distance of 0.033 and 0.011 on the sim-low (NGS) and sim-high (NGS) datasets, respectively. In comparison, other state-of-the-art tools obtain L2 distances ranging from 0.090 to 0.438 for sim-low and from 0.025 to 0.063 for sim-high. On PacBio HiFi data, PanTax attains L2 distances of 0.037 and 0.009 on the respective datasets, whereas alternative methods yield distances between 0.089 and 0.148 for sim-low and between 0.024 and 0.038 for sim-high. Similarly, on ONT R10.4 data, PanTax secures L2 distances of 0.03 and 0.009 for the datasets, while other tools achieve distances in the range of 0.09 to 0.148 for sim-low and 0.024 to 0.038 for sim-high. In addition, PanTax also consistently achieves the lowest absolute frequency error (AFE) and relative frequency error (RFE) across all sequencing data types (see Supplementary Table 1).

**Table 5.** Benchmarking results (L2 distance) of species-level taxonomic profiling on the simulated datasets (sim-low and sim-high). Note that the best score is marked in bold, and the second best score is marked in italics. N/A: not applicable.

| Methods | sim-low |  |  |  |  | sim-high |  |  |  |  |
| --- | --- | --- | --- | --- | --- | --- | --- | --- | --- | --- |
|  | NGS | PacBio | PacBio | ONT | ONT | NGS | PacBio | PacBio | ONT | ONT |
|  |  | HiFi | CLR | R9.4.1 | R10.4 |  | HiFi | CLR | R9.4.1 | R10.4 |
| PanTax | **0.033** | **0.037** | **0.017** | **0.018** | **0.030** | **0.011** | **0.009** | **0.009** | **0.008** | **0.009** |
| CLARK | 0.100 | 0.148 | 0.093 | 0.092 | 0.093 | 0.027 | *0.024* | 0.025 | 0.025 | 0.025 |
| KrakenUniq | 0.101 | 0.095 | 0.095 | 0.095 | 0.095 | 0.027 | 0.026 | 0.026 | 0.026 | 0.026 |
| Kraken | 0.160 | 0.148 | 0.148 | 0.148 | 0.148 | 0.039 | 0.038 | 0.038 | 0.038 | 0.038 |
| Kraken2 | 0.100 | 0.095 | 0.095 | 0.095 | 0.095 | 0.027 | 0.026 | 0.027 | 0.026 | 0.026 |
| Centrifuge | 0.099 | 0.091 | 0.090 | 0.091 | 0.091 | 0.027 | *0.024* | *0.024* | 0.024 | *0.024* |
| Bracken | 0.092 | N/A | N/A | N/A | N/A | *0.025* | N/A | N/A | N/A | N/A |
| MetaPhlAn4 | *0.090* | N/A | N/A | N/A | N/A | *0.025* | N/A | N/A | N/A | N/A |
| Qmatey | 0.438 | N/A | N/A | N/A | N/A | 0.063 | N/A | N/A | N/A | N/A |
| MetaMaps | N/A | 0.090 | 0.090 | *0.089* | *0.090* | N/A | *0.024* | 0.025 | 0.025 | *0.024* |
| MegaBLAST | N/A | *0.089* | *0.089* | *0.089* | *0.090* | N/A | *0.024* | *0.024* | 0.024 | 0.025 |

#### Pre-simulated datasets

As shown in Table 6, the benchmarking results for species-level taxonomic profiling across two pre-simulated datasets demonstrate varying performances among the tools. On the CAMI strain-madness dataset, PanTax achieves the second-best L2 distance on NGS data and the best L2 distance on PacBio CLR data; PanTax also achieves the second-best AFE/RFE on NGS data, and the best AFE and the second best RFE on PacBio CLR data (see Supplementary Table 2). In contrast, for the CAMI Gastrointestinal tract dataset, CLARK delivers the best performance (L2/AFE/RFE) on both NGS and PacBio data. Specifically, on the NGS data, PanTax achieves an L2 distance of 0.271, which is comparable to Centrifuge (0.249) and Qmatey (0.267), and significantly better than MetaPhlAn4 (0.421). On the PacBio data, while PanTax does not show a distinct advantage in terms of L2 distance, AFE and RFE, it still outperforms MetaMaps and MegaBLAST.

**Table 6.** Benchmarking results (L2 distance) of species-level taxonomic profiling on the pre-simulated datasets (CAMI strain-madness and CAMI Gastrointestinal tract). Note that the best score is marked in bold, and the second best score is marked in italics. N/A: not applicable.

| Methods | CAMI strain-madness |  | CAMI Gastrointestinal tract |  |
| --- | --- | --- | --- | --- |
|  | NGS | PacBio CLR | NGS | PacBio CLR |
| PanTax | *0.252* | **0.285** | 0.271 | 0.287 |
| CLARK | 0.355 | 0.389 | **0.239** | **0.171** |
| KrakenUniq | 0.354 | 0.389 | 0.243 | *0.174* |
| Kraken | 0.534 | 0.547 | 0.315 | 0.229 |
| Kraken2 | 0.356 | 0.379 | *0.240* | 0.203 |
| Centrifuge | 0.327 | 0.376 | 0.249 | 0.232 |
| Bracken | 0.386 | N/A | 0.306 | N/A |
| MetaPhlAn4 | **0.035** | N/A | 0.421 | N/A |
| Qmatey | 0.466 | N/A | 0.267 | N/A |
| MetaMaps | N/A | 0.400 | N/A | 0.333 |
| MegaBLAST | N/A | *0.371* | N/A | 0.320 |

#### Real datasets

As shown in Table 7, the benchmarking results for species-level taxonomic profiling across various real datasets demonstrate that PanTax achieves the best L2 distance on 4 out of 8 datasets, specifically Zymo1 (ONT R9.4.1/R10.4) and NWC (PacBio/ONT). It also secures the second-best L2 distance on two datasets: Zymo1 (NGS) and ATCC (PacBio HiFi). Notably, on the ATCC (PacBio HiFi) data, PanTax achieves the best AFE and RFE (see Supplementary Table 3). On the remaining two datasets, such as the ATCC (NGS) dataset, PanTax (0.199) has a worse L2 distance than MetaPhlAn4 (0.04) but achieves a comparable L2 distance to CLARK, Kraken2, KrakenUniq, and Centrifuge, and a better L2 distance than Kraken, Bracken, and Qmatey. Importantly, PanTax achieves the lowest RFE (0.320) while MetaPhlAn4 only achieves the second-best RFE (0.391) on this dataset. On the Zymo2 (HiFi) dataset, PanTax (0.291) exhibits slightly worse L2 distance in comparison with Centrifuge (0.263) and CLARK (0.268), and achieves comparable L2 when regarding MetaMaps (0.296), Kraken2 (0.299), and KrakenUniq (0.295). However, PanTax achieves the second-best RFE (0.610) on this dataset, which is slightly worse than MetaMaps (0.604) (see Supplementary Table 3).

**Table 7.** Benchmarking results (L2 distance) of species-level taxonomic profiling on the real datasets (Zymo1, Zymo2, ATCC, NWC). MegaBLAST was killed after running with 64 threads for more than fifteen days. Note that the best score is marked in bold, and the second best score is marked in italics. N/A: not applicable.

| Methods | Zymo1<br>NGS | Zymo1<br>ONT<br>R9.4.1 | Zymo1<br>ONT<br>R10.4 | Zymo2<br>HiFi | ATCC<br>NGS | ATCC<br>HiFi | NWC<br>PacBio | NWC<br>ONT |
| --- | --- | --- | --- | --- | --- | --- | --- | --- |
| PanTax | *0.308* | **0.241** | **0.272** | 0.291 | 0.199 | *0.228* | **0.247** | **0.364** |
| CLARK | 0.372 | *0.325* | *0.351* | *0.268* | 0.196 | **0.227** | 0.276 | 0.405 |
| KrakenUniq | 0.375 | 0.347 | 0.378 | 0.295 | 0.196 | 0.232 | 0.276 | 0.406 |
| Kraken | 0.376 | 0.349 | 0.382 | 0.341 | 0.308 | 0.293 | 0.401 | 0.513 |
| Kraken2 | 0.368 | 0.338 | 0.363 | 0.299 | 0.195 | *0.228* | 0.274 | 0.404 |
| Centrifuge | 0.357 | 0.327 | 0.354 | **0.263** | *0.192* | 0.237 | *0.273* | 0.402 |
| Bracken | 0.321 | N/A | N/A | N/A | 0.233 | N/A | N/A | N/A |
| MetaPhlAn4 | **0.193** | N/A | N/A | N/A | **0.040** | N/A | N/A | N/A |
| Qmatey | 0.613 | N/A | N/A | N/A | 0.435 | N/A | N/A | N/A |
| MetaMaps | N/A | 0.346 | 0.383 | 0.296 | N/A | 0.276 | 0.283 | *0.396* |
| MegaBLAST | N/A | 0.330 | 0.363 | - | N/A | - | 0.274 | 0.405 |

**Table 8.** Benchmarking results of strain-level taxonomic binning on the simulated datasets (sim-low and sim-high). AUPR: area under the precision-recall curve. Note that the best score is marked in bold, and the second best score is marked in italics.

| Methods | sim-low |  |  |  | sim-high |  |  |  |
| --- | --- | --- | --- | --- | --- | --- | --- | --- |
|  | Precision | Recall | F1 score | AUPR | Precision | Recall | F1 score | AUPR |
| *NGS* |  |  |  |  |  |  |  |  |
| PanTax | **0.949** | **0.933** | **0.941** | *0.916* | **0.812** | *0.781* | **0.796** | *0.685* |
| Centrifuge | *0.104* | **0.933** | *0.187* | **0.918** | *0.075* | **0.788** | *0.137* | **0.727** |
| Qmatey | 0.000 | 0.000 | 0.000 | 0.000 | 0.000 | 0.000 | 0.000 | 0.000 |
| PacBio HiFi |  |  |  |  |  |  |  |  |
| PanTax | **0.714** | *0.917* | **0.803** | 0.900 | **0.797** | *0.775* | **0.786** | *0.689* |
| Centrifuge | 0.549 | **0.933** | 0.691 | **0.930** | 0.298 | **0.787** | 0.432 | **0.739** |
| MetaMaps | *0.609* | **0.933** | *0.737* | *0.909* | *0.352* | 0.776 | *0.485* | 0.681 |
| PacBio CLR |  |  |  |  |  |  |  |  |
| PanTax | **0.636** | **0.933** | **0.757** | **0.926** | **0.734** | *0.778* | **0.755** | *0.696* |
| Centrifuge | 0.033 | **0.933** | 0.064 | *0.922* | 0.159 | **0.788** | 0.264 | **0.738** |
| MetaMaps | *0.548* | *0.850* | *0.667* | 0.688 | *0.374* | 0.770 | *0.503* | 0.550 |
| ONT R9.4.1 |  |  |  |  |  |  |  |  |
| PanTax | **0.691** | **0.933** | **0.794** | *0.909* | **0.755** | *0.781* | **0.768** | *0.701* |
| Centrifuge | 0.062 | **0.933** | 0.115 | **0.929** | 0.143 | **0.788** | 0.243 | **0.742** |
| MetaMaps | *0.448* | **0.933** | *0.605* | 0.780 | 0.236 | 0.777 | *0.363* | 0.579 |
| ONT R10.4 |  |  |  |  |  |  |  |  |
| PanTax | **0.709** | **0.933** | **0.806** | *0.912* | **0.776** | *0.779* | **0.777** | *0.687* |
| Centrifuge | 0.170 | **0.933** | 0.288 | **0.930** | 0.159 | **0.787** | 0.264 | **0.743** |
| MetaMaps | *0.533* | **0.933** | *0.679* | 0.851 | *0.227* | 0.777 | *0.352* | 0.624 |

### Strain level taxonomic binning for multiple species

Few methods currently available are suitable for strain-level taxonomic binning or profiling across multiple species (see Table 1). To ensure a fair comparison, we selected all methods capable of strain-level taxonomic classification for multiple species for our benchmarking evaluations. Specifically, we assessed Centrifuge, Qmatey, and CAMMiQ (which failed to run) on NGS data, and Centrifuge and MetaMaps on TGS data. Only the results of two freshly simulated datasets and one real dataset (Zymo1) are presented, as we cannot ascertain the truth of strain-level classification on pre-simulated and other real datasets.

#### Simulated datasets

On NGS data, PanTax (0.949/0.812) significantly outperforms Centrifuge (0.104/0.075) in terms of precision on both simulated datasets, while showing similar performance in recall and AUPR. Qmatey, however, did not yield any meaningful strain-level results. Comparable trends are seen in TGS data. For instance, on PacBio CLR data, PanTax (0.636/0.734) far exceeds Centrifuge (0.033/0.159) and MetaMaps (0.548/0.374) in precision for both the sim-low and sim-high datasets. On ONT R10.4 data, PanTax achieves precision scores of 0.709/0.776 across two datasets, whereas Centrifuge only achieves 0.170/0.159 and MetaMaps 0.533/0.227, respectively. Despite these differences in precision, PanTax generally maintains comparable recall to both Centrifuge and MetaMaps across nearly all TGS datasets. Its AUPR performance is slightly worse or on par with Centrifuge but substantially better than MetaMaps on most datasets.

#### Real datasets

To evaluate performance on the Zymo1 real datasets, we specifically reconstructed a new reference database so as to cover most of the strains in Zymo1. This reference database includes only eight species, but each species involves a greater number of strains selected from 34,697 complete genomes in RefSeq (as described in Step 1 of the Methods section). The number of strains per species is limited to 80 due to aligner performance constraints. Note that all benchmarking tools use this reference database for fair comparison. As shown in Table 10, the strain-level taxonomic binning results indicate that PanTax (0.875/0.5) significantly outperforms other methods (0.014/0.015) in terms of precision on both NGS and ONT data, while maintaining the same recall. Additionally, PanTax achieves the best AUPR on ONT data, though it performs less well on NGS data.

### Strain level taxonomic profiling for multiple species

#### Simulated datasets

Table 9 presents the results of strain-level taxonomic profiling across multiple species. For NGS data, PanTax (0.046/0.021) demonstrates superior performance in L2 distance compared to Centrifuge (0.109/0.026) and Qmatey (-/-) across both simulated datasets. On sim-low datasets (PacBio CLR, ONT R9.4.1/R10.4), PanTax also achieves the best L2 distance, but it performs worse than others on PacBio HiFi data. Conversely, on sim-high datasets (PacBio/ONT), PanTax achieves comparable L2 distance performance with the other methods. Furthermore, for NGS data, PanTax consistently outperforms other methods, achieving the best AFE and RFE scores on both datasets. However, for TGS data, PanTax’s AFE and RFE scores are largely on par with those of alternative approaches, as detailed in Supplementary Table 4.

**Table 9.**
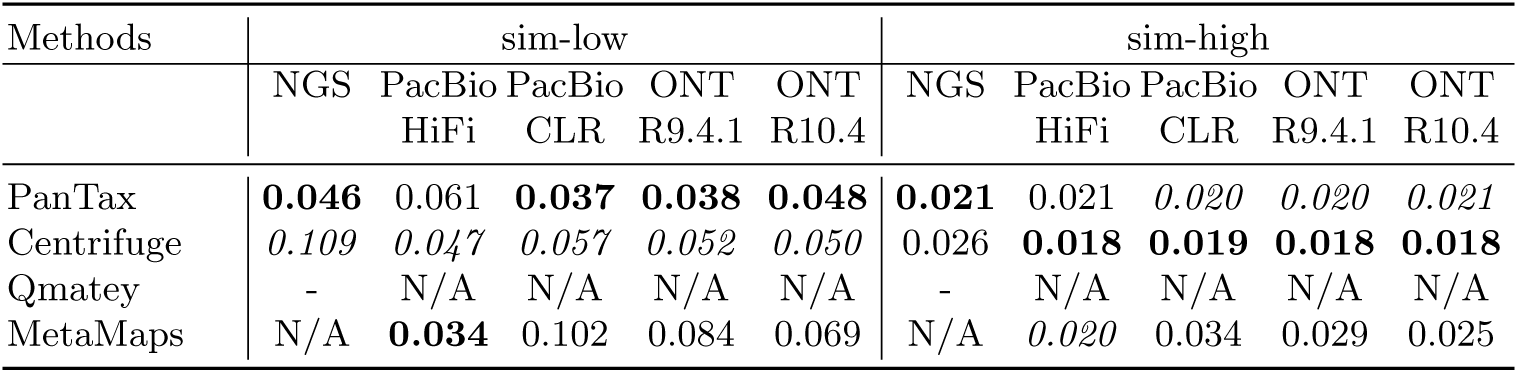
Benchmarking results (L2 distance) of strain-level taxonomic profiling on the simulated datasets (sim-low and sim-high). Qmatey did not return any matching strains when running on NGS dataset. Note that the best score is marked in bold, and the second best score is marked in italics. N/A: not applicable.

**Table 10.** Benchmarking results of strain-level taxonomic binning on the real datasets (Zymo1). Qmatey did not return any matching strains when running on NGS dataset. AUPR: area under the precision-recall curve. Note that the best score is marked in bold.

| Methods | Precision | Recall | F1 score | AUPR |
| --- | --- | --- | --- | --- |
| *NGS* |  |  |  |  |
| PanTax | **0.875** | **0.875** | **0.875** | 0.733 |
| Centrifuge | 0.014 | **0.875** | 0.027 | **0.858** |
| Qmatey | 0 | 0 | 0 | 0 |
| *ONT R9.4.1* |  |  |  |  |
| PanTax | **0.538** | **0.875** | **0.667** | **0.816** |
| Centrifuge | 0.013 | **0.875** | 0.026 | 0.789 |
| MetaMaps | 0.015 | **0.875** | 0.029 | 0.317 |
| *ONT R10.4* |  |  |  |  |
| PanTax | **0.500** | **0.875** | **0.636** | **0.748** |
| Centrifuge | 0.013 | **0.875** | 0.026 | 0.680 |
| MetaMaps | 0.015 | **0.875** | 0.029 | 0.262 |

#### Real datasets

Table 11 shows the results of strain-level taxonomic profiling on the Zymo1 datasets. PanTax demonstrates the lowest L2 distance across various data types, including NGS and ONT. In addition, PanTax significantly surpasses other methods in terms of AFE and RFE across both NGS and ONT sequencing data, as demonstrated in Supplementary Table 5.

**Table 11.** Benchmarking results (L2 distance) of strain-level taxonomic profiling on the real datasets (Zymo1). Note that the best score is marked in bold. N/A: not applicable.

| Methods | NGS | ONT<br>R9.4.1 | ONT<br>R10.4 |
| --- | --- | --- | --- |
| PanTax | **0.202** | **0.221** | **0.274** |
| Centrifuge | 0.494 | 0.250 | 0.311 |
| Qmatey | - | N/A | N/A |
| MetaMaps | N/A | 0.354 | 0.389 |

### Strain level taxonomic classification for single species

We note that several prominent metagenomic tools for strain-level taxonomic classification, such as Centrifuge and MetaMaps, have the capability to identify multiple species or strains and determine their relative abundances within a metagenomic sample. In addition to these versatile tools, there are others specifically designed to generate strain-level abundance profiles but are tailored for single-species analysis. These tools require users to preassign the species they wish to analyze, after which they provide outputs detailing the abundances of strains within the specified species. Noteworthy examples of such tools include StrainScan, StrainGE, and StrainEst (see Table 1).

To compare PanTax with other methods capable of performing strain-level microbiome composition analysis for single species from NGS reads, we conducted several benchmarking experiments. We chose *Staphylococcus epidermidis* (species taxid: 1282) as a example. When dealing with a single species, we chose all non-redundant genomes (70 strains) of this species using single-linkage clustering with the threshold of ANI≤99.9%. These genomes were subsequently utilized to construct the reference pangenome graph for this species. As shown in Table 12, PanTax achieved nearly perfect performance on all three datasets, attaining 100% precision, recall, and AUPR, as well as nearly zero L2 distance. This significantly outperforms other state-of-the-art approaches, such as StrainScan, StrainGE and StrainEst.

**Table 12.** Benchmarking results of strain-level composition analysis for single species using NGS data.We employed distinct strains of *Staphylococcus epidermidis* (genome size is about 2.5Mbp) in the aforementioned three datasets, with the pairwise ANI ranging from 99.2% to 99.3%, 98.6% to 99.3%, and 97.0% to 99.5%, respectively, across the three datasets.The methods are sorted in each metagenomic dataset by the precision in descending order.

| Methods | Precision<br>(strain) | Recall<br>(strain) | F1 score<br>(strain) | AUPR<br>(strain) | L2<br>distance | AFE | RFE |
| --- | --- | --- | --- | --- | --- | --- | --- |
| 3 strains |  |  |  |  |  |  |  |
| PanTax | **1.000** | **1.000** | **1.000** | **1.000** | **0.000** | **0.000** | **0.000** |
| StrainScan | **1.000** | 0.667 | 0.800 | 0.500 | 0.137 | 0.074 | 0.427 |
| StrainGE | **1.000** | **1.000** | **1.000** | **1.000** | 0.152 | 0.068 | 0.152 |
| StrainEst | 0.176 | **1.000** | 0.300 | **1.000** | 0.130 | 0.047 | 0.137 |
| 5 strains |  |  |  |  |  |  |  |
| PanTax | **1.000** | **1.000** | **1.000** | **1.000** | **0.000** | **0.000** | **0.000** |
| StrainScan | **1.000** | 0.800 | 0.889 | 0.700 | 0.058 | 0.019 | 0.287 |
| StrainGE | **1.000** | **1.000** | **1.000** | **1.000** | 0.136 | 0.050 | 0.221 |
| StrainEst | 0.294 | **1.000** | 0.455 | **1.000** | 0.071 | 0.020 | 0.155 |
| 10 strains |  |  |  |  |  |  |  |
| PanTax | **1.000** | **1.000** | **1.000** | **1.000** | **0.024** | **0.004** | **0.031** |
| StrainScan | **1.000** | 0.900 | 0.947 | 0.850 | 0.044 | 0.013 | 0.249 |
| StrainGE | **1.000** | 0.500 | 0.667 | 0.450 | 0.186 | 0.051 | 0.659 |
| StrainEst | 0.357 | **1.000** | 0.526 | **1.000** | 0.026 | 0.005 | 0.103 |

### Effects of divergence versus ratio of abundances

To explore the impact of divergence and abundance disparities on strain identification in mixed samples, we simulated multiple mixtures comprising two strains. Similarly, we used the reference database for *Staphylococcus epidermidis* (species taxid: 1282) from the previous section and selected 6 strains from it to perform the experiment. These mixtures encompassed various combinations of ANI values of 96.8%, 97%, 98%, 99%, and 99.8% (with 99.8% representing the most challenging scenario and 96.8% the least) and abundance ratios of 1:1, 1:3, 1:5, and 1:10. Our experiments centered on NGS reads, and we subsequently assessed performance of PanTax using F1 score, AUPR, and L2 distance for each possible combination of divergence and abundance ratio. The results are presented in Supplementary Figure 1. Analysis of the F1 score and AUPR indicates that PanTax successfully identifies both true strains in all scenarios. Regarding L2 distance, while PanTax accurately estimates the relative abundances of each strain in most cases, it exhibits biased abundance estimations for the most challenging scenario (ANI=99.8%).

### Effects of reducing sequencing coverage

To study the effects of reducing sequencing coverage, we randomly subsampled 1/2 and 1/10 of the reads from the “sim-low” datasets, naming these subsets “sim-low-sub1” and “sim-low-sub2”, respectively. We then benchmarked all related methods accordingly. The species-level taxonomic classification results can be found in Table 13 and Supplementary Table 6. Considering the low coverage strains in the dataset, we adjusted the threshold of *f*_strain_ to 0. These results show that PanTax consistently achieves superior precision, recall, F1 score, AUPR, AFE, and RFE across all sequencing data types, including NGS, PacBio HiFi/CLR, and ONT R9.4.1/R10.4, again at one single exception: for NGS data, PanTax achieves second-best precision. While MetaPhlAn4 attains higher species-level precision than PanTax, it suffers from significantly lower species-level recall. This discrepancy is particularly pronounced when sequencing coverage is ultra-low (e.g., median coverage less than 0.1x), as shown in Supplementary Table 8. The reason for this is quite intuitive: the marker based approach bases its assessment only on genome information that was proven to reliably distinguish between well-known species. However, it misses to identify species for which marker genes are not sufficiently available.

**Table 13.**
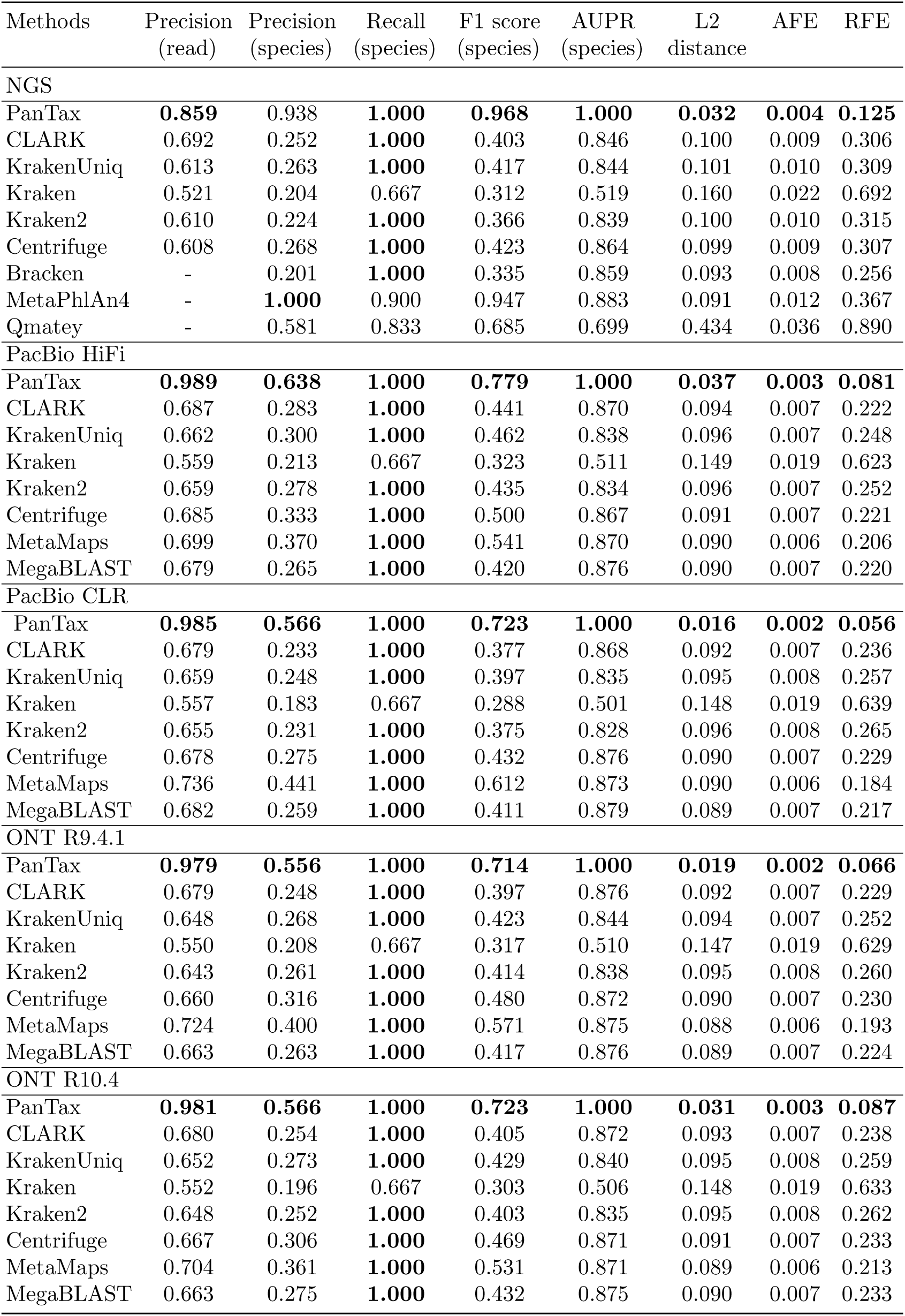
Benchmarking results of species-level taxonomic classification on the simulated dataset sim-low-sub1. The mean sequencing coverage of strains is about 2.65x. The 2nd column represents the result of individual read classification, whereas 4-10 columns represent the results of species. AFE: absolute frequency error, RFE: relative frequency error. Note that the best score is marked in bold.

In addition, refer to Table 14 and Supplementary Table 7 for strain-level taxonomic classification results on the sim-low-sub1 and sim-low-sub2 datasets, respectively. For sim-low-sub1, PanTax significantly outperforms other methods in terms of strain-level precision across all sequencing data types, including NGS, PacBio HiFi/CLR, and ONT R9.4.1/R10.4. It also maintains comparable or superior performance in metrics such as recall, AUPR, L2 distance, AFE and RFE. For sim-low-sub2, PanTax substantially outperforms Centrifuge in precision across all sequencing data types. While PanTax outperforms MetaMaps on ONT data, it shows slightly worse performance on PacBio data in terms of precision. PanTax also exhibits inferior recall and AUPR. In terms of L2 distance in taxonomic profiling, PanTax achieves better performance than Centrifuge on NGS data but worse performance on TGS data.

**Table 14.**
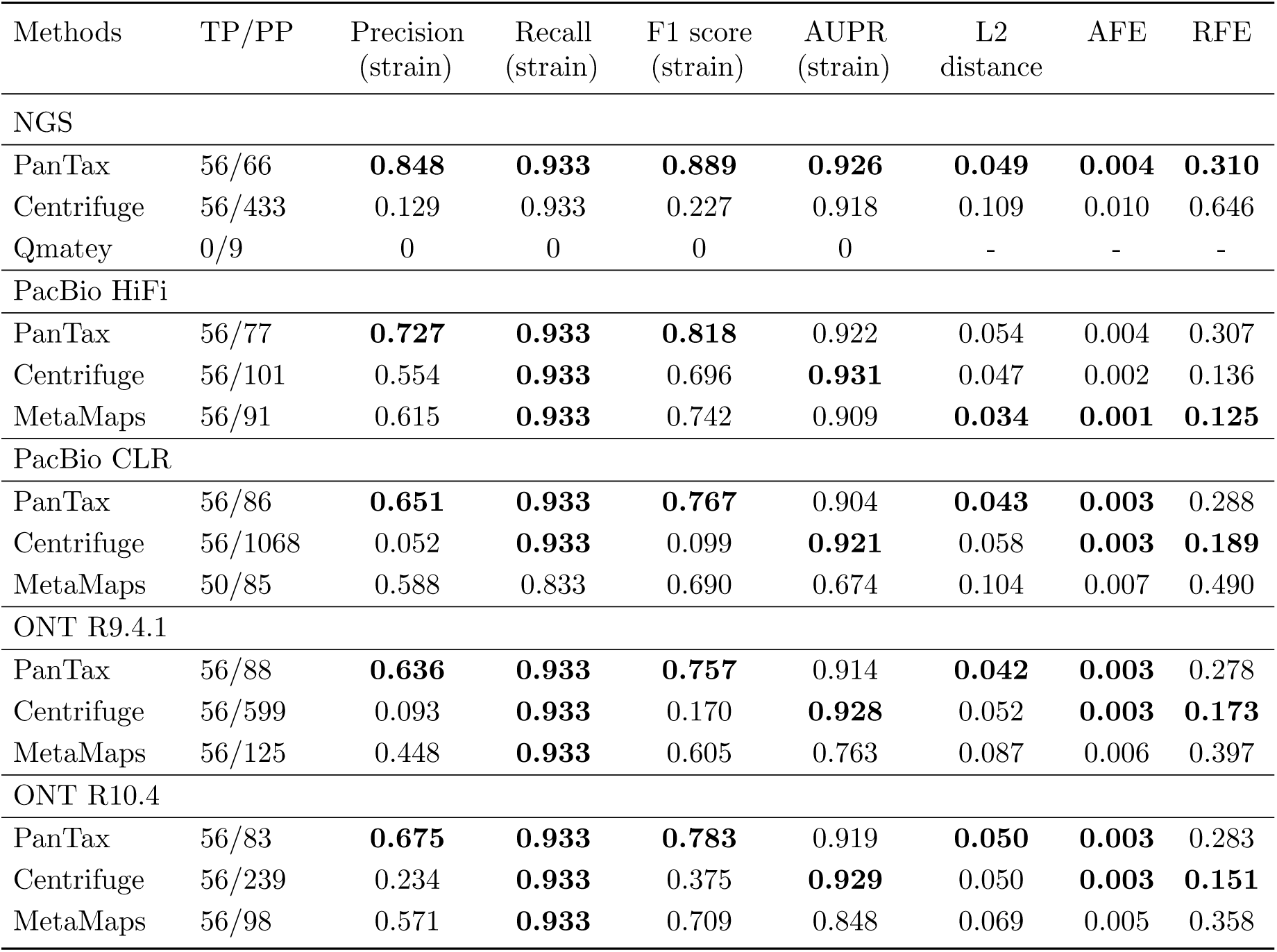
Benchmarking results of strain-level taxonomic classification on the simulated dataset sim-low-sub1. Note that Qmatey failed to predict any true strains. TP/PP: number of truly predicted positive strains (true positive) / number of predicted positive strains reported by methods (true positive + false positive), of which the value is equal to the precision in the third column.

| Methods | TP/PP | Precision<br>(strain) | Recall<br>(strain) | F1 score<br>(strain) | AUPR<br>(strain) | L2<br>distance | AFE | RFE |
| --- | --- | --- | --- | --- | --- | --- | --- | --- |
| NGS |  |  |  |  |  |  |  |  |
| PanTax | 56/66 | **0.848** | **0.933** | **0.889** | **0.926** | **0.049** | **0.004** | **0.310** |
| Centrifuge | 56/433 | 0.129 | 0.933 | 0.227 | 0.918 | 0.109 | 0.010 | 0.646 |
| Qmatey | 0/9 | 0 | 0 | 0 | 0 | - | - | - |
| PacBio HiFi |  |  |  |  |  |  |  |  |
| PanTax | 56/77 | **0.727** | **0.933** | **0.818** | 0.922 | 0.054 | 0.004 | 0.307 |
| Centrifuge | 56/101 | 0.554 | **0.933** | 0.696 | **0.931** | 0.047 | 0.002 | 0.136 |
| MetaMaps | 56/91 | 0.615 | **0.933** | 0.742 | 0.909 | **0.034** | **0.001** | **0.125** |
| PacBio CLR |  |  |  |  |  |  |  |  |
| PanTax | 56/86 | **0.651** | **0.933** | **0.767** | 0.904 | **0.043** | **0.003** | 0.288 |
| Centrifuge | 56/1068 | 0.052 | **0.933** | 0.099 | **0.921** | 0.058 | **0.003** | **0.189** |
| MetaMaps | 50/85 | 0.588 | 0.833 | 0.690 | 0.674 | 0.104 | 0.007 | 0.490 |
| ONT R9.4.1 |  |  |  |  |  |  |  |  |
| PanTax | 56/88 | **0.636** | **0.933** | **0.757** | 0.914 | **0.042** | **0.003** | 0.278 |
| Centrifuge | 56/599 | 0.093 | **0.933** | 0.170 | **0.928** | 0.052 | **0.003** | **0.173** |
| MetaMaps | 56/125 | 0.448 | **0.933** | 0.605 | 0.763 | 0.087 | 0.006 | 0.397 |
| ONT R10.4 |  |  |  |  |  |  |  |  |
| PanTax | 56/83 | **0.675** | **0.933** | **0.783** | 0.919 | **0.050** | **0.003** | 0.283 |
| Centrifuge | 56/239 | 0.234 | **0.933** | 0.375 | **0.929** | 0.050 | **0.003** | **0.151** |
| MetaMaps | 56/98 | 0.571 | **0.933** | 0.709 | 0.848 | 0.069 | 0.005 | 0.358 |

In summary, these experiments demonstrate that PanTax is the superior choice of approach in particular when dealing with low coverage (components of) metagenomes.

### Direct comparison of pangenome graphs and multiple linear genomes

For species-level taxonomic classification, although most existing classifiers that can handle multiple species typically select a single representative genome as the reference for each species, it is also possible to use multiple representative genomes for classification by manually adapting the reference database. To compare the performance of PanTax, using pangenome graphs, with other tools, using multiple linear genomes, we conducted benchmarking experiments on the sim-low (NGS) and CAMI strain-madness (NGS) datasets. All benchmarking methods utilized RefDB:13404 as the reference database, employing either graph-based or linear representations. As shown in Supplementary Table 9, PanTax achieves superior species-level precision, F1 score, and L2 distance while maintaining comparable species-level recall and AUPR on both datasets compared to other methods that use multiple linear genomes as references. Overall, this provides clear evidence of the superiority of pangenome graph-based reference systems over using multiple linear reference genomes for taxonomic classification.

### Runtime and memory usage evaluation

Supplementary Table 10 presents the runtime and peak memory usage of benchmarking tools utilizing NGS data. We primarily compare the runtime and memory usage for index construction and read alignment during queries, as these processes dominate the main computational resources. When dealing with NGS reads, PanTax initially requires index construction for the pangenome graphs, whereas this indexing process is unnecessary for TGS reads. Notably, the indexing process accounts for the majority of the time compared to the read-to-graph alignment. However, for a given pangenome reference, indexing needs to be performed only once. PanTax exhibits a shorter index construction time compared to other tools, including Kraken, Kraken2, KrakenUniq, and Centrifuge. During the classification step, PanTax is slightly slower than Kraken2 and CLARK, yet it achieves comparable efficiency with Centrifuge. It is worth noting that for large datasets, such as “sim-high”, PanTax still consumes approximately three times less overall CPU time than MetaPhlAn4. Additionally, PanTax demands slightly higher peak memory than other methods, namely Kraken, KrakenUniq, CLARK, and Centrifuge. Supplementary Table 11 demonstrates that, on most TGS datasets, PanTax exhibits a slower runtime compared to other methods, being several times slower. It also demands a higher peak memory compared to the alternative methods.

## Discussion

We have introduced PanTax, as an approach that classifies the contents of metagenomes in terms of their taxa. PanTax accepts all classes of popular sequencing reads as input, and determines the organisms that make part of the metagenome not only at the level of species, but also at the level of strains. Apart from one exception, PanTax is the only approach that gathers all essential qualities required for comprehensive assessment of metagenomes in terms of taxonomic classification of their contents: there is no bias with respect ot pre-selected genomic regions, it can process all popular types of reads, it operates at the finest level of taxonomy (i.e. at the level of strains), it can handle multiple species simultaneously, it can integrate custom databases—which ensures to keep up with the already fast space in terms of the detection of novel bacterial species and strains—it provides estimates on the abundances of the different taxa, and it can classify individual reads. Experiments on simulated and real data demonstrate that PanTax substantiallyy outperforms the only prior approach that gathers all of these qualities. By and large, PanTax also outperforms or is on a par with state-of-the-art methods that specialize in particular aspects of taxonomic classification of metagenoms.

Key to success is the fact that PanTax, to the best of our knowledge, is the first approach that builds on pangenome graphs as reference systems, instead of linear genomes. Unlike ordinary linear reference genomes, pangenome graphs establish reference systems that can capture the diversity inherent to a collection of genomes even at the level of strains, the finest taxonomic resolution possible. The type of pangenome graph that PanTax is based on are variation graphs. Beyond just capturing the diversity of a mix of genomes, variation graphs also arrange all genomes in an evolutionarily consistent manner, which eliminates ambiguities during classification.

This lack of ambiguities is particularly beneficial when determining the strain or species specific origin of single reads: because a read aligns with a graph that incorporates all genomes evolutionarily consistently, the read-to-graph alignment immediately points out the strain (or species) the read stems from. Instead, aligning a read against the applicable linear reference genomes entails a statistically involved analysis of the alignment scores that each of the different alignments delivers; because the single linear genomes do not explicitly relate with each other, this analysis may leave one with various statistical uncertainties. A particularly interesting scenario results from failure of a read to align with any of the linear reference genomes. Still, however, the read may nevertheless align with the graph that has integrated all the genomes the read does not satisfyingly align with. A situation like this indicates that the read aligns with a hitherto unknown strain of the species represented by the graph, and means that the read stems from that species, which one may overlook when operating with linear genomes alone.

Further, additional advantages of pangenome graphs are due to the compact representation of related genomes. First, the compactness of the graphs renders the execution of alignments simply more efficient: instead of having to align a read with each of the linear genomes separately, one read-to-graph alignment suffices. Secondly, the read-to-graph mappings are not only less ambiguous, but also quite simply more accurate, in particular in the presence of structural variations (Garrison *et al*., 2018; Siŕen *et al*., 2021). Last but not least, variation graphs, as the (arguably most popular) type of pangenome graphs in use here, facilitate the formulation of an optimization problem that we have termed “path abundance optimization problem”, which gives rise to strain-level taxonomic classification in a tractable way. This optimization problem based strategy allows for utmost nuanced and accurate distinction between the strains of a species. In summary, read-to-pangenome-graph alignments do not only offer a unified approach to align a read with many related sequences, as above-mentioned, but they also offer a way that is unified with respect to the particular type of reads that one deals with. They can be reliably applied for both accurate short and noisy long reads (as well as reads that are both accurate and long, of course), which k-mer based alignment-free strategies cannot guarantee.

Benchmarking experiments have demonstrated that the improvements raised by PanTax relative to key metrics for evaluating taxonomic classification are substantial, if not, at times, even drastic. Note that we sought select experiments such that they were maximally diverse in terms of types of metagenome sequencing reads used and alternative approaches compared with. While PanTax outperformed all state-of-the-art approaches from an overall point of view, this does not mean that selected prior approaches did not achieve competitive performance rates, or even slightly outperformed PanTax on certain selected combinations of evaluation categories. For example, MetaPhlAn4 rivalled PanTax in terms of classifying accurate, short NGS reads at the level of species; for example, MetaPhlAn4 achieved better precision, while PanTax achieved better recall. In turn, however, PanTax outperforms MetaPhlAn4 at the level of species.

Importantly, PanTax tends to outperform other approaches even in their favorable categories when decreasing read volumes to ultra-low coverage. This delivers decisive arguments that PanTax both captures low-abundance strains at greater accuracy and why lowering expenses in terms of read volume generation is justified when using PanTax. In the light of the ongoing developments in terms of read sequencing technologies, it is further convincing to see that PanTax outperforms all other approaches in terms of species-level classification on the great majority of datasets when receiving TGS data as input. The advantages of PanTax span across all of PacBio HiFi (accurate), PacBio CLR (noisy), ONT R10.4 and ONT R9.4.1, which provides evidence of its unequivocal advantages when exploiting technological advances in terms of sequencing.

Whole-genome based methods(e.g. PanTax) generally offer greater sensitivity and overall performance metrics, such as Recall and AUPR, by leveraging extensive genomic data. In contrast, marker-based methods(e.g. MetaPhlAn4) excel in precision for species-level identification due to their focus on specific, well-characterized genomic markers. However, these methods face limitations in identifying species lacking known markers, distinguishing strains, and processing third-generation sequencing data. Additionally, the marker-based approach can be slower due to the complexity of marker gene identification.

Another look at the comparison of PanTax with the leading marker based approach MetaPhlAn4 points out that the whole-genome based approach PanTax excels in terms of Recall and AUPR while the marker based approach achieves better Precision in species-level binning and L2 distance in species-level profiling. The reason for this is quite intuitive: the marker based approach bases its assessment only on genome information that was proven to reliably distinguish between well-known species. However, it misses to identify species for which marker genes are not sufficiently available. Furthermore, marker-based approaches fail to distinguish between strains and classify individual sequencing reads, in addition to missing support for TGS data (because no longer undergoing regular extensions) and being very slow due to the expenses in terms of the identification of marker genes.

We have also demonstrated the superiority of utilizing pangenome graph based representations of multiple genomes over simply making use of multiple linear genomes that one, for example, concatenates to one long sequence for obtaining a naive, sequence based representation of multiple linear genomes. The corresponding experiments referring to an immediate comparison of pangenome graph with multiple linear genomes provide conclusive evidence that much more than anything else the pangenome *graphs* are the decisive, advantageous factor in the task of taxonomic classification. Note that advantages persist on increasing numbers of genomes, which points out that the number of genomes integrated in either graphs or used as linear genomes is fairly irrelevant: pangenome graphs have decisive advantages in all cases. Strain-level classification poses an even greater methodical challenge, because sequential differences between strains can look near-negligible at first glance. So, strain-level analyses of metagenomes have only recently gained pace. This explains why there are only few existing approaches, and why all of the are very recent. PanTax outperforms all other approaches that aim at classifying metagenomes at the level of strains quite significantly. PanTax does so for both NGS and TGS data, across the entire board of evaluation metrics: it is the only approach that outperforms others significantly, while preserving competitive performance rates also in the categories where other approaches have their particular advantages. Last but not least, PanTax proves to outperform the other approaches in strain-level classification of isolate species; although isolates are not the main intention of our approach, these experiments demonstrate that PanTax is not negatively affected by scenarios characterized by a few dominant species—on the contrary.

In summary, PanTax is compatible with NGS, both noisy PacBio CLR and accurate PacBio HiFi, as well as ONT, both noisy R9.4 and the more accurate R10.4 (error rate about 2%) sequencing data. It supports classification at both the species and the strain level, for both single and multiple species, so establishes an approach that is universal with respect to all current scenarios of interest in metagenomics type experiments. While Centrifuge is the only currently existing method that rivals PanTax in terms of universality, PanTax outperforms Centrifuge on the vast majority of benchmark categories quite substantially, so proves to be the preferable universal classifier. We recall that, unlike PanTax, Centrifuge is not based on pagenome graphs. This underscores the superiority of pangenome graph based taxonomic classification another time.

Further improvements of PanTax are conceivable and they are promising. Currently, just like any other reference based classification approach, PanTax is unable to generate taxonomic profiles for strains that, because they are novel, are missing in the reference databases. Instead, PanTax reports the most similar strain recorded in the databases. While this at least accurately determines the most likely closest relative, a preferable solution would be to dynamically update the species-specific pangenome graphs by incorporating hitherto unobserved genetic variation. From a larger perspective, this would support the detection of unknown strains, beyond just enhancing classification by removing mistaken hits.

Improvements in terms of runtimes are also conceivable. Because PanTax is less trimmed towards efficient usage of computational resources than other methods, PanTax tends to consume more runtime than other approaches when working with larger databases. Solutions are already on the horizon: in the future, we plan on seed based strategies, known to substantially speed up the alignment of reads to sequences or graphs at virtually no losses in terms of the accuracy of the alignments.

## Methods

In the following, we provide the full range of methodical details involved in the steps in Figure 1. We continue with additional, relevant remarks regarding benchmark competitors, and we conclude by defining the metrics used for evaluating results.

### Pangenome graph based reference database construction

When constructing our default pangenome based reference database (“Graph representation of reference databases” in Fig. 1), we focus on bacteria as the primary object of interest in the majority of metagenomics studies. We are aware that in metagenomics studies also viruses, archaea, fungi, or plasmids (and so on) can be in the major focus of attention. To account for this, one can, mutatis mutandis, readily extend our methodology to other organisms. Users can flexibly feed their customized databases as input to PanTax, instead of using the provided default databases. Once databases are provided, everything else follows the identical workflow.

In the following, numbers and names of ‘Steps’ correspond to the numbering and naming of procedures in Figure 1.

#### Step 1: Select high-quality genomes for species

We retrieved high-quality, complete (in particular gap-free) bacterial genomes from NCBI’s RefSeq (O’Leary *et al*., 2016) on July 18, 2023 (RefSeq Release 219). The NCBI’s RefSeq (release 219) establishes a comprehensive reference database, consisting of 34,697 complete genomes. In view of plasmid sequences being typically considerably shorter than bacterial genomes, plasmid sequences were excluded from our experiments. We remind that one can put particular attention also to plasmids, if desirable, by augmenting our database with plasmid reference sequence based pangenomes, for example. For each species, we computed a pairwise distance matrix using FastANI (Jain *et al*., 2018) where rows and columns correspond to the strains of the species under consideration, and single entries in the matrix indicate the ANI of two strains of that species. Subsequently, we applied a customized graph-based clustering approach (see Algorithm 1 in the Supplementary Methods) at ANI thresholds of 95% and 99.9%, ensuring that the mutual ANI’s of the strain-specific genomes for each species fell within this range. To maintain a balance between sensitivity and computational cost, we determined no more than 10 representative genomes for each species as a reasonable choice. Following these guidelines, we included 8,778 species, amounting to a total of 13,404 strains, in the refined reference database. Of the 8,778 species included, 1,313 comprised multiple strains (see Supplementary Figure 2).

#### Step 2: Construct a pangenome graph for each species

Subsequently, we constructed a variation graph (as the particular, and generally most popular type of pangenome graph) for each species, by using the (efficient and unbiased) pangenome graph builder PGGB (Garrison *et al*., 2023). We construct the variation graph *G* = (*V, E, P*) from the (multiple) strain genomes pertaining to a species, where *V* are the nodes, *E* are the edges and *P* are the paths in *G* corresponding to the input strain-specific genomes used for constructing the graph. *G* is a bidirected sequence graph, where each vertex *v* ∈ *V* corresponds to a sequence seq(*v*) that consists of single or multiple nucleotides (A, T, C, G).

Edges *e_ij_* = (*v_i_, v_j_*) ∈ *E* indicate that the sequences seq(*v_i_*) and seq(*v_j_*) represented by nodes *v_i_*and *v_j_* appear as a sequential subsequence in one of the genomes that contribute to the graph. Because paths *P* outline these genomes, paths *P* cover the graph, or, in other words, each edge is contained in one of the paths from *P*. Let 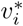 denote the reverse-complement of node *v_i_*. Then the reverse edge 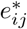 of *e_ij_* is defined as 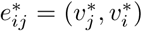. Both nodes and edges can be traversed in either the forward or the reverse direction in the graph. In general, known strains can be identified as the specific paths *p* = (*v*_1_*, …, v_l_*) ∈ *P*, as part of the definition of the species-specific pangenome graph *G*.

For species that involve only one strain, the task is to turn its corresponding sequence into a dedicated variation graph, acting as representative counterpart to the pangenome graphs constructed from multiple strains. Here, we achieved this by partitioning the genome sequence into fragments of (the relatively short) length 1024bp, each of which is to correspond to a node in the graph, which, by design, is chain-like.

Subsequently, one merges all species-specific pangenome graphs into a single, large pangenome graph, by relabeling nodes, while never introducing new nodes or edges. Of note, the species-specific pangenome graphs can be computed in paraellel, which avoids computational bottlenecks in this step. As outlined above, this procedure supports the unified and unambiguous classification of single reads via *one* read-to-graph alignment, instead of having to align a read with each of the (here, by default: 8,778) species-specific pangenome graphs individually. In other words, instead of having to perform and evaluate 8,778 alignments in statistically involved mutual context, the merging of graphs delivers one encompassing, optimal and statistically sensible alignment that directly indicates the species the read optimally aligns with.

### Species-level taxonomic classification

#### Step 3: Sequence to graph alignment

Sequencing reads are then mapped to the merged pangenome graph using existing, approved and efficient sequence-to-graph aligners. For short reads, we employ Giraffe (Siŕen *et al*., 2021), as an integral part of the vg toolkit (Garrison *et al*., 2018) supporting the computation of sequence-to-graph alignments for preferably short reads. For long reads, we make use GraphAligner (Rautiainen and Marschall, 2020) by default.

For faster alignment of long reads, we also have implemented an option (--fast) that utilizes the Giraffe aligner in its long-read mode instead of GraphAligner. Note that the long read mode of Giraffe is currently under development such that alignments tend to be worse than those computed with GraphAligner. Despite the current disadvantages of Giraffe when using long reads, we have nevertheless implemented it, in expectation of future progress thanks to the rapid developments it is currently undergoing.

The basis for speeding up read alignment with Giraffe is the construction of a graph Burrows-Wheeler transform (GBWT) (Siŕen *et al*., 2020) which one uses for indexing the pangenome graph. For PanTax, we resort to building the GBWT index manually, via a ‘fast indexing mode’, which relies on three commands provided by vg, which considerably speeds up index construction. Although the default fully automatic construction of the index via ‘vg autoindex’ is supposed to be more accurate, classification results did not differ in our experiments, which explains our choice.

#### Step 4: Species-level taxonomic binning

Taxonomic binning involves the allocation of sequencing reads to distinct taxonomic groups. In this procedure, we assign each mapped read to a species-specific pangenome graph, utilizing the results of the sequence-to-graph alignment resulting from aligning reads with the merged pangenome graph. When a read aligns to multiple graphs (within this unifying merged graph), we only retain the optimal alignment, thereby ensuring that each read corresponds to a single species. We again note that the optimal alignment is easy to determine precisely because the merging of graphs implies that different alignment scores can be put into mutual context.

Subsequently, we assess whether individual species need to be flagged as false positives. That is, although the species receives a high score, the species does not make part of the metagenome. For that, we examine the MAPQ (mapping quality) scores of all the reads whose optimal alignments correspond to the particular species. We raise two key criteria: first, we require at least one read to reach a MAPQ of 60, which indicates a statistically significant unique match of at least one read with that species. We further require that at least one tenth of the reads that optimally align with a species exhibit a MAPQ of at least 2; if less than one tenth of the reads aligned with a species have MAPQ less than 2, the species is flagged as false positive. It remains to mention that reads may fail to align with the pangenome graph, which potentially indicates the presence of novel strains (not necessarily from novel species, although that is possible as well) in the metagenomic sample under analysis. We discard such unmapped reads from further consideration in the subsequent analysis.

#### Step 5: Species-level taxonomic profiling

For species *i*, we divide the total base count of the reads that optimally aligned with species *i* via the read-to-graph alignment by the average length of the strain-specific genomes that contributed to constructing the pangenome graph for species *i*, which we refer to as *normalized read coverage c_i_* in the following. Subsequently, we calculate the relative abundance of species *i* as the ratio 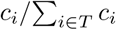, where *T* is the set of species reported in the previous step (e.g. resulting from excluding potentially false positives when dealing with NGS data, before outputting all species referred to as *T* here).

### Strain-level taxonomic classification

#### Step 6: Path abundance optimization

Furthermore, we offer an optional strain-level classification. Having already constructed a species-specific pangenome graph by collapsing identical sequences from multiple strains (as described in Step 2), we formulate a path abundance optimization problem for each species-specific pangenome graph, following an approach suggested in (Baaijens *et al*., 2020). In this step, the abundance we refer to is absolute abundance, which is the average coverage depth. This step aims to resolve strain-level taxonomic classification. Let *V* be the set of all nodes in a species-specific pangenome graph, *P* be the set of paths corresponding to all strains used to construct the pangenome graph for a particular species (where a strain’s genome sequence corresponds to a path in the pangenome graph in general), and 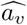 be the absolute abundance of node *v* ∈ *V* in the graph, as estimated by evaluating the reads whose alignments include *v* (i.e., the average coverage depth of a single base in this node), where we follow the definitions provided in (Baaijens *et al*., 2020) in a one-to-one fashion. For a path *p* ∈ *P* (again following (Baaijens *et al*., 2020)), we define *a_p_* ∈ R_≥0_ to be the absolute abundance of path *p*, which corresponds to the abundance of the strain that corresponds to *p*. Here, we would like to predict all *a_p_*by raising them as variables in the program

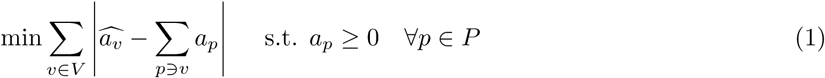

which we refer to as *path abundance optimization (PAO) problem*. Values *a_p_* ≥ 0 establishing a solution are output as the abundance values of the strains that correspond to the paths *p*.

As for an explanation, the objective function of the PAO problem aims to estimate the abundance (i.e. the normalized read coverage) of each strain by minimizing the difference between the abundances of the nodes one has observed (derived from the sequence-to-graph alignments in Step 3) and the abundances of the nodes that correspond to the abundances of the strains whose genomes include the nodes. The PAO problem is convex in nature, and can be efficiently linearized and solved using a linear programming solver of choice; here we opted for Gurobi (Gurobi Optimization, 2023). So, from a theoretical perspective, this problem is polynomial-time solvable. In addition, since the size of the candidate set of strain paths *P* is typically not overly large (usually ≤ 10), the optimization problems can be solved in parallel across the species, which renders this step computationally very efficient.

Note that for each species, we conduct two iterations of path abundance optimization. To further mitigate the odds of raising false positives during classification, we add a filtering operation prior to each iteration of PAO. To illustrate the filtering step, we first define a *triplet node* as a set of three consecutive nodes within a path. A triplet node that is exclusive to a single strain, that is the three nodes only appear in that order in one of the paths *p* ∈ *P*, is referred to as *strain-specific triplet node*.

(1) For the filtering step that precedes the first iteration of solving the PAO, let *V*_reads_ be the consecutive node triples covered by reads aligning with the species-specific pangenome graph and *V*_strain_ reflect the set of strain-specific triplet nodes that show in the path corresponding to the strain, as just defined. We then raise the metric f, as per the definition

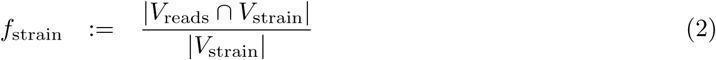

As per its definition, *f* captures the proportion of strain-specific triplet nodes in the strain genome (reference) the concatenation of the three sequences of which appears in the reads of the sequencing sample. As a ratio, the value of *f* naturally ranges from 0 to 1, with 1 indicating that all strain-specific triplet nodes are covered by reads from the sample. We then filter out strains where *f*_strain_ *<* 0.3.

Subsequently, we solve the PAO problem a first time for only the strains that were not discarded in the filtering procedure (that is whose *f*_strain_ was found to be ≥ 0.3), and record the strains, respectively the paths *p* in the species-specific pangenome graph to which they correspond whose *a_p_ >* 0, that is we keep all strains that make part of the solution of the PAO.

(2) For the filtering step that precedes the second iteration of solving the PAO, let **v**_triplet_ refer to triplet nodes, as defined above, and let *c*(**v**_triplet_) be the average read coverage of **v**_triplet_, where averaging is across nucleotides in the sequence of **v**_triplet_. Further, we introduce *a_p,_*_triplet_, which, unlike *a_p_* that resulted from the solution of the PAO, estimates the abundance of the strain that corresponds to *p* by taking the average of the coverages of the triplet nodes, that is the average of the *c*(**v**_triplet_) in *p*:

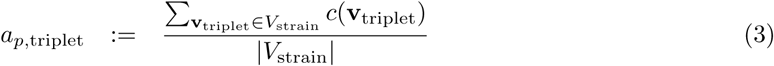

Based on this defintion, we raise

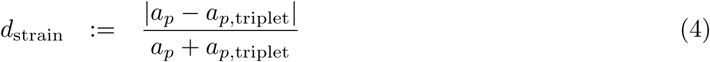

As for a brief explnation, *d*_strain_ quantifies the the divergence between the strain abundance *a_p_* predicted in the first iteration of the PAO and the strain abundance estimate based on the average coverage of all strain-specific triplet nodes.

Before proceeding with the second iteration of the PAO, we filter out further potentially false positive strains by keeping only strains where *d*_strain_ ≤ 0.45.

Finally, we solve the PAO a second time. The final output of strains are the ones whose *a_p_* is greater than zero in that second iteration.

#### Step 7: Strain-level taxonomic profiling

The final, additional step is to provide relative abundance estimates for any strain predicted to make part of the sample. For that, one determines the absolute abundances of the strains, and divides them by the absolute abundance of the species they belong to, as predicted in step 5. There is one caveat: the sum of the absolute abundances of the strains could exceed the absolute abundance of their species. In that case, we scale the absolute abundances of the strains such that their sum equals that of their species. The relative abundance of one strain then is the appropriately normalized / scaled version of the absolute abundance of the strain, as just determined.

### Metrics for evaluation

We evaluate the performance of taxonomic classifiers using metrics that are widely adopted in various relevant studies (Simon *et al*., 2019; Zhu *et al*., 2022; Luo *et al*., 2022a). The primary criteria for assessing metagenomic classification methods are precision and recall. These metrics can be computed at the level of individual reads (task: assigning reads to species or strains), and also at the levels of species and strains (task: determine which species or strains make part of the sample). Precision, at the read classification level, quantifies the fraction of correctly classified reads in the sample over all reads that are classified (i.e. assigned to a genome) by the method. At the level of taxa (species and strains), Precision measures the fraction of correctly identified taxa among all taxa that were identified. Recall, at the level of taxa, evaluates the proportion of correctly identified taxa compared to all taxa present in the sample. It is important to note that we refrain from offering the read-level recall metric, as formulating a reasonable and consistent definition for such a measure poses significant challenges.

The F1 score, the harmonic mean of precision and recall, is frequently employed as a balanced metric for evaluating classifiers.

However, precision, recall, and the F1 score may not reflect the performance of classifiers in metagenomic samples sufficiently comprehensively, because the F1 score neglects the correct assessment of low-abundance taxa, whose accurate detection, however, can be crucial. A approach to evaluating the contents of metagenomes that takes low-abundance taxa into sufficiently accurate account is to evaluate the metagenomes in terms of curves that plot precision and recall across varying abundance thresholds. Calculating the area under this curve (“AUPR”) yields a metric that does not neglect the performance of a classifier also with respect to taxa of lower abundance (Simon *et al*., 2019). For the calculation of AUPR, refer to (Simon *et al*., 2019). Classifiers that fail to recall all ground-truth taxa are penalized with a zero AUPR score from their greatest recall achieved up to 100% recall. Conversely, classifiers that achieve 100% recall are not further penalized in their AUPR score for additional false positive taxon calls. Please see “AUPR calculation example” in the Supplementary Methods for illustrative examples.

Beyond the accurate identification of taxa only, it can be imperative to provide accurate estimates of the abundances of the taxa that make part of a metagenome. An example for the necessity of such practice are metagenome-wide association studies (“MetaGWAS”) where fluctuations in the proportions of the participating taxa can have profound implications in terms of the phenotypes supported by the metagenome. The relative abundances of taxa across various samples are comparable, making them a primary focus in MetaGWAS studies. The ultimate output of taxonomic profiling at the species or strain level (Steps 5 or 7) is precisely these relative abundances. Notably, in this study, we propose a set of metrics to assess the accuracy and reliability of these relative abundance estimates. To determine abundance profiles for tools that can output read assignment, we follow the exact same protocol employed for PanTax. Namely, we determine species level abundances for the alternative methods as per Step 5 (see above). There is one difference, when scaling coverages with respect to genome length, the length of the (single) representative genome that the respective method employs is used, which reflects the consistent equivalent for linear genome based methods. However, PanTax employs the average length of all strain genomes within a species to normalize read coverages, as it utilizes a pangenome based approach rather than relying solely on a single linear genome. If methods offer abundance profiling as output directly, we make use of that. Note that for the species level binning, only species with relative abundance greater than 0.0001, following the recommendations provided in Zhu *et al*. (2022) are used without special circumstances. The L2 distance, a commonly used metric, serves as a quantitative evaluation of abundance profiles (Simon *et al*., 2019). Given a sample, let *I* represent the intersection set of estimated and true taxa. For a taxon *i* ∈ *I*, let 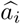 and *a_i_* represent the estimated and true abundance of taxon *i*, respectively. Then, the L2 distance is computed as follows: 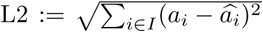. A smaller L2 distance indicates a higher similarity between the estimated and true abundances, while greater L2 distance suggests greater divergence. However, it is worth noting that the L2 distance is particularly sensitive to the accurate quantification of highly abundant taxa within a sample, as these taxa exert a significant influence on the overall similarity between abundance profiles. To address this, we introduce two complementary metrics: the absolute frequency error (AFE) and the relative frequency error (RFE). AFE is defined as: 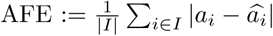, whereas RFE is defined as: 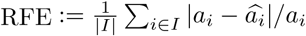. These two metrics have been adapted from previous studies on viral haplotype abundance evaluation (Luo *et al*., 2022a; Baaijens *et al*., 2020). The AFE and RFE metrics provide insights into the deviation of estimated abundances from their true values. Notably, the RFE metric balances the influence of low-abundance species or strains, because it scales divergences relative to the true abundance *a_i_* of the taxa.

## Data availability

The genomes and simulated sequencing reads generated by this study can be downloaded from Zenodo DOI: https://doi.org/10.5281/zenodo.11221154.

## Code availability

The source code of PanTax is GPL-3.0 licensed, and publicly available at https://github.com/LuoGroup2023/PanTax. The results presented in this study can be reproduced from Code Ocean under DOI: https://codeocean.com/capsule/6917504/tree/v1.

## Acknowledgements

XL is supported by the National Natural Science Foundation of China (Grant No. 32400506), the Natural Science Foundation of Hunan Province (Grant No. 2024JJ4008) and Fundamental Research Funds for the Central Universities (Grant No. 541109030062). AS received funding from the European Union’s Horizon 2020 research and innovation programme under Marie Sk-lodowska-Curie grant agreements No 956229 (ALPACA) and No 872539 (PANGAIA). YL is supported by the National Natural Science Foundation of China (Grant No. 62372159).

## Author contributions

Xiao Luo conceived this study. Xiao Luo, Wenhai Zhang and Yuansheng Liu designed the method. Wenhai Zhang implemented the software. Xiao Luo and Alexander Schönhuth wrote and revised the manuscript. Wenhai Zhang, Yuansheng Liu, Jialu Xu and Enlian Chen conducted the data analysis. All authors read and approved the final version of the manuscript.

