## Supplementary Material for "PanTax: Strain-level taxonomic classification of metagenomic data using pangenome graphs"

#### Supplementary Tables and Figures

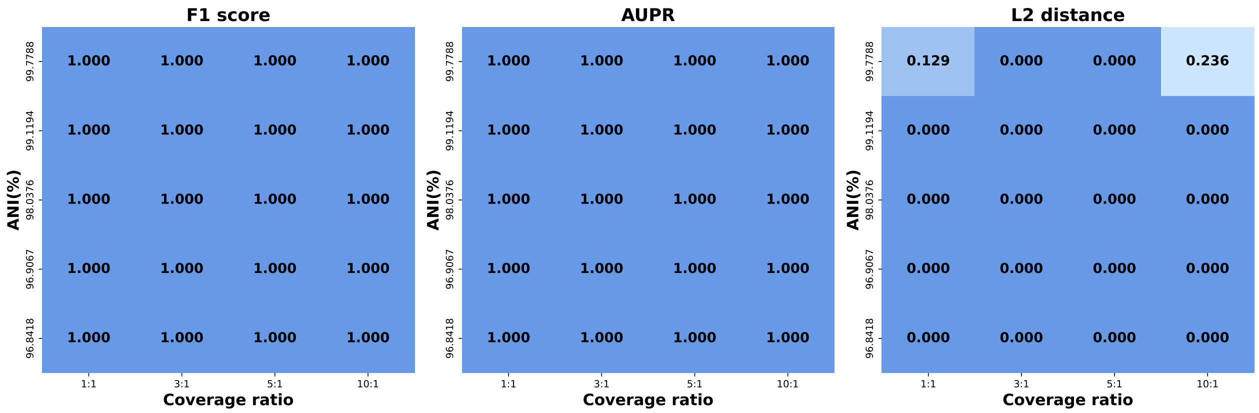


**Supplementary Figure 1.** Evaluation of PanTax's performance in mixtures of two strains (of the same species) with varying strain coverage and ANI using simulated NGS reads. The x-axis represents the coverage ratio, while the y-axis depicts the ANI (Average Nucleotide Identity) of the two strains in the mixture. The three panels correspond to the F1 score, AUPR (Area Under the Precision-Recall Curve), and L2 distance of the strain-level taxonomic classification results, respectively. Darker shades indicate superior taxonomic performance.

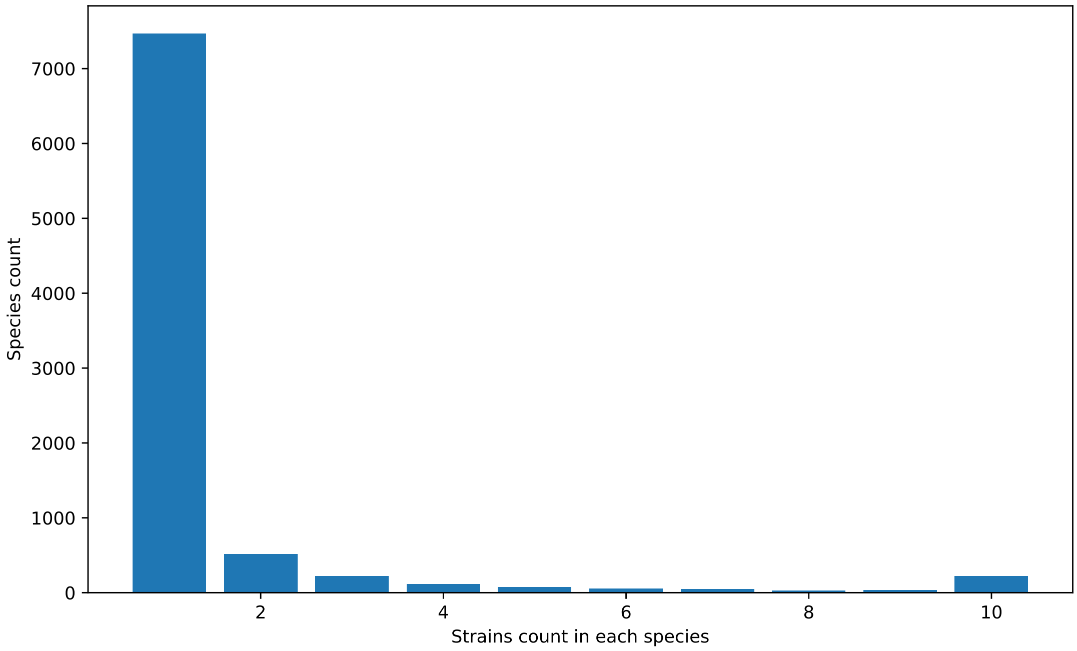


**Supplementary Figure 2.** Distribution of strain numbers across species. The bar chart depicts the number of strains (genomes) represented for each species in the pangenome reference dataset used in this study. The X-axis indicates the number of strains within each species, while the Y-axis represents the number of species.

| Methods | sim-low | | | sim-high | | |
|---|---|---|---|---|---|---|
| | Precision (read) | AFE | RFE | Precision (read) | AFE | RFE |
| *NGS* | | | | | | |
| PanTax | **0.859** | **0.004** | **0.125** | **0.880** | **0.000** | **0.115** |
| CLARK | 0.692 | 0.009 | 0.306 | 0.793 | 0.001 | 0.292 |
| KrakenUniq | 0.613 | 0.010 | 0.308 | 0.663 | 0.001 | 0.297 |
| Kraken | 0.521 | 0.022 | 0.693 | 0.520 | 0.002 | 0.570 |
| Kraken2 | 0.610 | 0.010 | 0.315 | 0.658 | 0.001 | 0.297 |
| Centrifuge | 0.609 | 0.009 | 0.307 | 0.652 | 0.001 | 0.296 |
| Bracken | - | 0.008 | 0.256 | - | 0.001 | 0.231 |
| MetaPhlAn4 | - | 0.012 | 0.359 | - | 0.001 | 0.272 |
| Qmatey | - | 0.012 | 0.359 | - | 0.002 | 0.854 |
| PacBio HiFi | | | | | | |
| PanTax | **0.970** | **0.003** | **0.067** | **0.944** | **0.000** | **0.061** |
| CLARK | 0.690 | 0.007 | 0.216 | 0.768 | 0.001 | 0.193 |
| KrakenUniq | 0.666 | 0.007 | 0.242 | 0.745 | 0.001 | 0.207 |
| Kraken | 0.563 | 0.019 | 0.627 | 0.576 | 0.001 | 0.488 |
| Kraken2 | 0.662 | 0.007 | 0.245 | 0.742 | 0.001 | 0.209 |
| Centrifuge | 0.687 | 0.007 | 0.215 | 0.759 | 0.001 | 0.192 |
| MetaMaps | 0.704 | 0.006 | 0.199 | 0.782 | **0.000** | 0.178 |
| MegaBLAST | 0.681 | 0.007 | 0.214 | 0.743 | 0.001 | 0.204 |
| PacBio CLR | | | | | | |
| PanTax | **0.972** | **0.002** | **0.052** | **0.940** | **0.000** | **0.060** |
| CLARK | 0.679 | 0.007 | 0.229 | 0.756 | 0.001 | 0.204 |
| KrakenUniq | 0.659 | 0.007 | 0.251 | 0.734 | 0.001 | 0.214 |
| Kraken | 0.556 | 0.019 | 0.640 | 0.569 | 0.001 | 0.498 |
| Kraken2 | 0.655 | 0.008 | 0.258 | 0.727 | 0.001 | 0.218 |
| Centrifuge | 0.678 | 0.007 | 0.223 | 0.745 | 0.001 | 0.200 |
| MetaMaps | 0.737 | 0.006 | 0.182 | 0.807 | **0.000** | 0.175 |
| MegaBLAST | 0.682 | 0.007 | 0.214 | 0.746 | 0.001 | 0.202 |
| ONT R9.4.1 | | | | | | |
| PanTax | **0.979** | **0.002** | **0.055** | **0.944** | **0.000** | **0.060** |
| CLARK | 0.679 | 0.007 | 0.223 | 0.761 | 0.001 | 0.200 |
| KrakenUniq | 0.648 | 0.007 | 0.247 | 0.714 | 0.001 | 0.212 |
| Kraken | 0.549 | 0.019 | 0.633 | 0.554 | 0.001 | 0.493 |
| Kraken2 | 0.643 | 0.007 | 0.252 | 0.707 | 0.001 | 0.215 |
| Centrifuge | 0.660 | 0.007 | 0.223 | 0.722 | 0.001 | 0.198 |
| MetaMaps | 0.721 | 0.006 | 0.191 | 0.788 | **0.000** | 0.177 |
| MegaBLAST | 0.664 | 0.007 | 0.217 | 0.739 | 0.001 | 0.202 |
| ONT R10.4 | | | | | | |
| PanTax | **0.960** | **0.003** | **0.072** | **0.932** | **0.000** | **0.070** |
| CLARK | 0.679 | 0.007 | 0.226 | 0.759 | 0.001 | 0.201 |
| KrakenUniq | 0.651 | 0.007 | 0.249 | 0.721 | 0.001 | 0.212 |
| Kraken | 0.551 | 0.019 | 0.633 | 0.559 | 0.001 | 0.491 |
| Kraken2 | 0.647 | 0.007 | 0.251 | 0.716 | 0.001 | 0.214 |
| Centrifuge | 0.666 | 0.007 | 0.222 | 0.730 | 0.001 | 0.198 |
| MetaMaps | 0.703 | 0.006 | 0.201 | 0.776 | 0.001 | 0.183 |
| MegaBLAST | 0.662 | 0.007 | 0.221 | 0.736 | 0.001 | 0.206 |

**Supplementary Table 1.** Benchmarking results of species-level read binning and taxonomic profiling on the simulated datasets (sim-low and sim-high). Bracken, MetaPhlAn4 and Qmatey do not return read alignment. Note that the best score is marked in bold.

| Methods | CAMI strain-madness | | | CAMI Gastrointestinal tract | | |
|---|---|---|---|---|---|---|
| | Precision (read) | AFE | RFE | Precision (read) | AFE | RFE |
| *NGS* | | | | | | |
| PanTax | **0.411** | 0.019 | 0.711 | 0.461 | 0.024 | 0.645 |
| CLARK | 0.339 | 0.033 | 1.468 | **0.537** | **0.021** | **0.488** |
| KrakenUniq | 0.189 | 0.032 | 1.503 | 0.434 | **0.021** | 0.490 |
| Kraken | 0.076 | 0.049 | 0.907 | 0.336 | 0.024 | 0.625 |
| Kraken2 | 0.184 | 0.033 | 1.440 | 0.433 | **0.021** | 0.501 |
| Centrifuge | 0.221 | 0.031 | 1.542 | 0.435 | 0.022 | 0.525 |
| Bracken | - | 0.036 | 1.112 | - | 0.027 | 0.641 |
| MetaPhlAn4 | - | **0.005** | **0.661** | - | **0.021** | 0.522 |
| Qmatey | - | 0.048 | 1.290 | - | 0.023 | 0.603 |
| PacBio CLR | | | | | | |
| PanTax | **0.545** | **0.028** | 1.613 | 0.493 | 0.024 | 0.565 |
| CLARK | 0.499 | 0.037 | 2.955 | **0.630** | **0.014** | **0.385** |
| KrakenUniq | 0.217 | 0.037 | 2.967 | 0.409 | **0.014** | 0.390 |
| Kraken | 0.075 | 0.049 | **0.917** | 0.309 | 0.017 | 0.531 |
| Kraken2 | 0.186 | 0.037 | 2.618 | 0.275 | 0.019 | 0.504 |
| Centrifuge | 0.198 | 0.039 | 2.591 | 0.301 | 0.022 | 0.544 |
| MetaMaps | 0.322 | 0.034 | 1.803 | 0.500 | 0.027 | 0.630 |
| MegaBLAST | 0.335 | 0.037 | 2.250 | 0.465 | 0.028 | 0.652 |

**Supplementary Table 2.** Benchmarking results of species-level read binning and taxonomic profiling on the pre-simulated datasets (CAMI strain-madness and CAMI Gastrointestinal tract). Bracken, MetaPhlAn4 and Qmatey do not return read alignment. Note that the best score is marked in bold.

| Methods | AFE | RFE | AFE | RFE |
|---|---|---|---|---|
| | Zymo1 NGS | | Zymo1 ONT R9.4.1 | |
| PanTax | 0.090 | 0.722 | **0.065** | **0.520** |
| CLARK | 0.111 | 0.890 | 0.091 | 0.725 |
| KrakenUniq | 0.112 | 0.898 | 0.094 | 0.752 |
| Kraken | 0.113 | 0.902 | 0.094 | 0.754 |
| Kraken2 | 0.110 | 0.881 | 0.093 | 0.741 |
| Centrifuge | 0.107 | 0.857 | 0.089 | 0.711 |
| Bracken | 0.086 | 0.689 | N/A | N/A |
| MetaPhlAn4 | **0.051** | **0.428** | N/A | N/A |
| Qmatey | 0.154 | 1.234 | N/A | N/A |
| MetaMaps | N/A | N/A | 0.082 | 0.654 |
| MegaBLAST | N/A | N/A | 0.085 | 0.677 |
| | Zymo2 HiFi | | ONT R10.4 | |
| PanTax | 0.050 | 0.610 | **0.075** | **0.597** |
| CLARK | **0.048** | 0.642 | 0.093 | 0.743 |
| KrakenUniq | 0.052 | 0.781 | 0.101 | 0.808 |
| Kraken | 0.061 | 0.854 | 0.102 | 0.814 |
| Kraken2 | 0.052 | 0.781 | 0.097 | 0.777 |
| Centrifuge | **0.048** | 0.651 | 0.091 | 0.730 |
| MetaMaps | 0.050 | **0.604** | 0.091 | 0.724 |
| MegaBLAST | - | - | 0.086 | 0.692 |
| | ATCC NGS | | ATCC HiFi | |
| PanTax | 0.019 | **0.320** | **0.020** | **0.496** |
| CLARK | 0.019 | 0.419 | 0.022 | 0.583 |
| KrakenUniq | 0.019 | 0.422 | 0.022 | 0.614 |
| Kraken | 0.036 | 0.632 | 0.033 | 0.785 |
| Kraken2 | 0.018 | 0.423 | 0.022 | 0.605 |
| Centrifuge | 0.018 | 0.417 | 0.022 | 0.614 |
| Bracken | 0.016 | 0.392 | N/A | N/A |
| MetaPhlAn4 | **0.004** | 0.391 | N/A | N/A |
| Qmatey | 0.050 | 1.475 | N/A | N/A |
| MetaMaps | N/A | N/A | 0.023 | 0.580 |
| MegaBLAST | N/A | N/A | - | - |
| | NWC PacBio | | NWC ONT | |
| PanTax | **0.133** | **0.473** | **0.199** | **0.677** |
| CLARK | 0.151 | 0.526 | 0.220 | 0.747 |
| KrakenUniq | 0.151 | 0.526 | 0.221 | 0.747 |
| Kraken | 0.203 | 0.760 | 0.277 | 0.986 |
| Kraken2 | 0.150 | 0.524 | 0.220 | 0.745 |
| Centrifuge | 0.152 | 0.523 | 0.220 | 0.741 |
| MetaMaps | 0.152 | 0.537 | 0.215 | 0.733 |
| MegaBLAST | 0.151 | 0.524 | 0.222 | 0.746 |

**Supplementary Table 3.** Benchmarking results of species-level taxonomic profiling on the real datasets (ATCC, Zymo1, Zymo2, NWC). MegaBLAST was killed after running for more than fifteen days. Note that the best score is marked in bold. N/A: not applicable.

| Methods | sim-low | | sim-high | |
|---|---|---|---|---|
| | AFE | RFE | AFE | RFE |
| NGS | | | | |
| PanTax | **0.003** | **0.183** | **0.000** | **0.350** |
| Centrifuge | 0.010 | 0.646 | 0.001 | 0.647 |
| Qmatey | - | - | - | - |
| PacBio HiFi | | | | |
| PanTax | 0.003 | 0.198 | **0.000** | 0.363 |
| Centrifuge | 0.002 | 0.124 | **0.000** | 0.284 |
| MetaMaps | **0.001** | **0.100** | **0.000** | **0.301** |
| PacBio CLR | | | | |
| PanTax | **0.002** | 0.184 | **0.000** | **0.343** |
| Centrifuge | 0.003 | **0.168** | **0.000** | 0.355 |
| MetaMaps | 0.007 | 0.472 | 0.001 | 0.628 |
| ONT R9.4.1 | | | | |
| PanTax | **0.002** | 0.172 | **0.000** | 0.340 |
| Centrifuge | 0.003 | **0.154** | **0.000** | **0.322** |
| MetaMaps | 0.005 | 0.346 | 0.001 | 0.533 |
| ONT R10.4 | | | | |
| PanTax | 0.003 | 0.186 | **0.000** | 0.353 |
| Centrifuge | **0.002** | **0.139** | **0.000** | **0.316** |
| MetaMaps | 0.006 | 0.387 | **0.000** | 0.446 |

**Supplementary Table 4.** Benchmarking results of strain-level taxonomic profiling on the simulated datasets (sim-low and sim-high). Qmatey failed to predict any true strains. Note that the best score is marked in bold.

| Methods | AFE | RFE |
|---|---|---|
| NGS | | |
| PanTax | **0.042** | **0.338** |
| Centrifuge | 0.128 | 1.025 |
| Qmatey | - | - |
| ONT R9.4.1 | | |
| PanTax | **0.056** | **0.447** |
| Centrifuge | 0.078 | 0.627 |
| MetaMaps | 0.104 | 0.834 |
| ONT R10.4 | | |
| PanTax | **0.076** | **0.609** |
| Centrifuge | 0.095 | 0.757 |
| MetaMaps | 0.112 | 0.893 |

**Supplementary Table 5.** Benchmarking results of strain-level taxonomic profiling on the real datasets (Zymo1). Qmatey failed to predict any true strains. Note that the best score is marked in bold.

| Methods | Precision (read) | Precision (species) | Recall (species) | F1 score (species) | AUPR (species) | L2 distance | AFE | RFE |
|---|---|---|---|---|---|---|---|---|
| NGS | | | | | | | | |
| PanTax | **0.859** | 0.938 | **1.000** | **0.968** | **1.000** | **0.033** | **0.004** | **0.125** |
| CLARK | 0.692 | 0.256 | **1.000** | 0.408 | 0.848 | 0.100 | 0.009 | 0.305 |
| KrakenUniq | 0.612 | 0.265 | **1.000** | 0.420 | 0.843 | 0.101 | 0.009 | 0.308 |
| Kraken | 0.520 | 0.198 | 0.667 | 0.305 | 0.519 | 0.159 | 0.022 | 0.690 |
| Kraken2 | 0.609 | 0.229 | **1.000** | 0.373 | 0.840 | 0.100 | 0.010 | 0.314 |
| Centrifuge | 0.608 | 0.265 | **1.000** | 0.420 | 0.866 | 0.099 | 0.009 | 0.307 |
| Bracken | - | 0.208 | 1.000 | 0.345 | 0.860 | 0.092 | 0.008 | 0.252 |
| MetaPhlAn4 | - | **1.000** | 0.867 | 0.929 | 0.883 | 0.100 | 0.014 | 0.426 |
| Qmatey | - | 0.690 | 0.667 | 0.678 | 0.572 | 0.421 | 0.037 | 0.938 |
| PacBio HiFi | | | | | | | | |
| PanTax | **0.990** | **0.625** | **1.000** | **0.769** | **1.000** | **0.038** | **0.003** | **0.105** |
| CLARK | 0.685 | 0.268 | **1.000** | 0.423 | 0.865 | 0.094 | 0.007 | 0.242 |
| KrakenUniq | 0.660 | 0.286 | **1.000** | 0.444 | 0.834 | 0.097 | 0.008 | 0.269 |
| Kraken | 0.562 | 0.217 | 0.667 | 0.328 | 0.508 | 0.150 | 0.019 | 0.620 |
| Kraken2 | 0.657 | 0.275 | **1.000** | 0.432 | 0.828 | 0.097 | 0.008 | 0.273 |
| Centrifuge | 0.686 | 0.312 | **1.000** | 0.476 | 0.862 | 0.092 | 0.007 | 0.242 |
| MetaMaps | 0.695 | 0.361 | **1.000** | 0.531 | 0.869 | 0.091 | 0.007 | 0.226 |
| MegaBLAST | 0.676 | 0.256 | **1.000** | 0.408 | 0.872 | 0.091 | 0.007 | 0.245 |
| PacBio CLR | | | | | | | | |
| PanTax | **0.985** | **0.566** | **1.000** | **0.723** | **1.000** | **0.020** | **0.003** | **0.096** |
| CLARK | 0.681 | 0.191 | **1.000** | 0.321 | 0.864 | 0.093 | 0.008 | 0.271 |
| KrakenUniq | 0.662 | 0.268 | **1.000** | 0.423 | 0.841 | 0.095 | 0.008 | 0.286 |
| Kraken | 0.557 | 0.189 | 0.667 | 0.294 | 0.504 | 0.147 | 0.019 | 0.642 |
| Kraken2 | 0.657 | 0.234 | **1.000** | 0.380 | 0.827 | 0.095 | 0.008 | 0.290 |
| Centrifuge | 0.678 | 0.268 | **1.000** | 0.423 | 0.876 | 0.090 | 0.007 | 0.260 |
| MetaMaps | 0.741 | 0.429 | **1.000** | 0.600 | 0.877 | 0.088 | 0.006 | 0.221 |
| MegaBLAST | 0.688 | 0.280 | **1.000** | 0.438 | 0.879 | 0.089 | 0.007 | 0.250 |
| ONT R9.4.1 | | | | | | | | |
| PanTax | **0.979** | **0.566** | **1.000** | **0.723** | **1.000** | **0.028** | **0.003** | **0.125** |
| CLARK | 0.672 | 0.259 | **1.000** | 0.411 | 0.875 | 0.093 | 0.008 | 0.257 |
| KrakenUniq | 0.642 | 0.270 | **1.000** | 0.426 | 0.848 | 0.096 | 0.008 | 0.282 |
| Kraken | 0.546 | 0.204 | 0.667 | 0.312 | 0.518 | 0.147 | 0.019 | 0.625 |
| Kraken2 | 0.635 | 0.268 | **1.000** | 0.423 | 0.833 | 0.097 | 0.009 | 0.292 |
| Centrifuge | 0.655 | 0.291 | **1.000** | 0.451 | 0.876 | 0.091 | 0.008 | 0.261 |
| MetaMaps | 0.720 | 0.390 | **1.000** | 0.561 | 0.880 | 0.088 | 0.007 | 0.220 |
| MegaBLAST | 0.656 | 0.270 | **1.000** | 0.426 | 0.868 | 0.091 | 0.008 | 0.261 |
| ONT R10.4 | | | | | | | | |
| PanTax | **0.981** | **0.545** | **1.000** | **0.706** | **1.000** | **0.036** | **0.004** | **0.131** |
| CLARK | 0.677 | 0.261 | **1.000** | 0.414 | 0.867 | 0.095 | 0.008 | 0.288 |
| KrakenUniq | 0.650 | 0.275 | **1.000** | 0.432 | 0.837 | 0.096 | 0.009 | 0.305 |
| Kraken | 0.551 | 0.211 | 0.667 | 0.320 | 0.502 | 0.150 | 0.020 | 0.646 |
| Kraken2 | 0.647 | 0.248 | **1.000** | 0.397 | 0.835 | 0.096 | 0.009 | 0.308 |
| Centrifuge | 0.664 | 0.297 | **1.000** | 0.458 | 0.874 | 0.092 | 0.008 | 0.280 |
| MetaMaps | 0.697 | 0.361 | **1.000** | 0.531 | 0.864 | 0.091 | 0.008 | 0.265 |
| MegaBLAST | 0.659 | 0.265 | **1.000** | 0.420 | 0.867 | 0.092 | 0.008 | 0.286 |

**Supplementary Table 6.** Benchmarking results of species-level taxonomic classification on the simulated dataset sim-low-sub2. The mean sequencing coverage of strains is about 0.53x. The 2nd column represents the result of individual read classification, whereas 3-10 columns represent the results of species. AUPR: area under the precision-recall curve, AFE: absolute frequency error, RFE: relative frequency error. Note that the best score is marked in bold.

| Methods | TP/PP | Precision (strain) | Recall (strain) | F1 score (strain) | AUPR (strain) | L2 distance | AFE | RFE |
|---|---|---|---|---|---|---|---|---|
| NGS | | | | | | | | |
| PanTax | 42/48 | **0.875** | 0.700 | **0.778** | 0.661 | **0.088** | **0.008** | 0.693 |
| Centrifuge | 56/274 | 0.204 | **0.933** | 0.335 | **0.919** | 0.109 | 0.010 | **0.645** |
| Qmatey | 0/6 | 0 | 0 | 0 | 0 | - | - | - |
| PacBio HiFi | | | | | | | | |
| PanTax | 42/60 | 0.700 | 0.700 | 0.700 | 0.668 | 0.093 | 0.009 | 0.744 |
| Centrifuge | 56/92 | 0.609 | **0.933** | 0.737 | **0.923** | 0.048 | 0.003 | 0.195 |
| MetaMaps | 56/77 | **0.727** | **0.933** | **0.818** | 0.910 | **0.035** | **0.002** | **0.166** |
| PacBio CLR | | | | | | | | |
| PanTax | 39/62 | 0.629 | 0.650 | 0.639 | 0.600 | 0.088 | 0.008 | 0.766 |
| Centrifuge | 56/355 | 0.158 | **0.933** | 0.270 | **0.918** | **0.059** | **0.003** | **0.235** |
| MetaMaps | 47/74 | **0.635** | 0.783 | **0.701** | 0.593 | 0.111 | 0.007 | 0.522 |
| ONT R9.4.1 | | | | | | | | |
| PanTax | 42/67 | **0.627** | 0.700 | **0.661** | 0.695 | 0.087 | 0.009 | 0.770 |
| Centrifuge | 56/246 | 0.228 | **0.933** | 0.366 | **0.930** | **0.056** | **0.003** | **0.236** |
| MetaMaps | 54/106 | 0.509 | 0.900 | 0.651 | 0.737 | 0.094 | 0.006 | 0.439 |
| ONT R10.4 | | | | | | | | |
| PanTax | 42/68 | **0.618** | 0.700 | 0.656 | 0.639 | 0.088 | 0.008 | 0.683 |
| Centrifuge | 56/142 | 0.394 | **0.933** | 0.554 | **0.927** | **0.053** | **0.003** | **0.194** |
| MetaMaps | 56/97 | 0.577 | **0.933** | **0.713** | 0.832 | 0.072 | 0.005 | 0.399 |

**Supplementary Table 7.** Benchmarking results of strain-level taxonomic classification on the simulated dataset sim-low-sub2. Note that Qmatey failed to predict any true strains. TP/PP: number of truly predicted positive strains (true positive) / number of predicted positive strains reported by methods (true positive + false positive), of which the value is equal to the precision in the third column. AUPR: area under the precision-recall curve.

| Methods | Precision (read) | Precision (species) | Recall (species) | F1 score (species) | AUPR (species) | L2 distance | AFE | RFE |
|---|---|---|---|---|---|---|---|---|
| NGS | | | | | | | | |
| PanTax | **0.745** | **0.964** | 0.900 | **0.931** | **0.880** | **0.204** | **0.013** | 0.360 |
| CLARK | 0.442 | 0.022 | **0.933** | 0.044 | 0.414 | 0.355 | 0.018 | **0.332** |
| KrakenUniq | 0.369 | 0.086 | **0.933** | 0.157 | 0.433 | 0.360 | 0.019 | 0.335 |
| Kraken | 0.349 | 0.065 | 0.600 | 0.117 | 0.269 | 0.464 | 0.027 | 0.727 |
| Kraken2 | 0.371 | 0.023 | **0.933** | 0.046 | 0.419 | 0.355 | 0.018 | 0.340 |
| Centrifuge | 0.369 | 0.020 | 0.900 | 0.038 | 0.451 | 0.356 | 0.019 | 0.354 |
| Bracken | - | 0.110 | 0.767 | 0.192 | 0.368 | 0.325 | 0.014 | 0.404 |
| MetaPhlAn4 | - | 1.000 | 0.433 | 0.605 | 0.417 | 0.214 | 0.015 | 0.746 |
| Qmatey | - | 0.625 | 0.167 | 0.263 | 0.070 | 0.503 | 0.026 | 0.925 |

**Supplementary Table 8.** Benchmarking results of species-level taxonomic classification using ultra-low coverage data. To demonstrate the limitations of MetaPhlAn4 in handling ultra-low coverage data, we simulated a new NGS dataset from scratch, which comprises 30 bacterial species, with one strain per species, and varying coverage levels distributed across six tiers: 5 species each at 18.0x, 1.8x, 0.18x, 0.018x, 0.0018x and 0.0002x average coverage levels, respectively. Considering the presence of ultra-low abundance species, the threshold abundance of species reported on this dataset was set to 0 instead of 0.0001.

| Methods | Precision (read) | Precision (species) | Recall (species) | F1 score (species) | AUPR (species) | L2 distance |
|---|---|---|---|---|---|---|
| sim-low | | | | | | |
| PanTax | 0.859 | **0.938** | **1.000** | **0.968** | **1.000** | **0.033** |
| CLARK | **0.996** | 0.857 | **1.000** | 0.923 | **1.000** | 0.064 |
| KrakenUniq | 0.728 | 0.882 | **1.000** | 0.938 | **1.000** | 0.064 |
| Kraken2 | 0.736 | 0.423 | **1.000** | 0.594 | 0.998 | 0.061 |
| Centrifuge | 0.731 | 0.750 | **1.000** | 0.857 | **1.000** | 0.062 |
| Bracken | - | 0.345 | **1.000** | 0.513 | 0.962 | 0.055 |
| CAMI strain-madness | | | | | | |
| PanTax | 0.411 | **0.135** | **0.750** | **0.229** | 0.335 | **0.252** |
| CLARK | **0.530** | 0.065 | **0.750** | 0.120 | 0.320 | 0.291 |
| KrakenUniq | 0.292 | 0.079 | **0.750** | 0.143 | 0.322 | 0.292 |
| Kraken2 | 0.291 | 0.068 | **0.750** | 0.125 | 0.315 | 0.287 |
| Centrifuge | 0.364 | 0.125 | **0.750** | 0.214 | **0.373** | 0.257 |
| Bracken | - | 0.063 | **0.750** | 0.116 | 0.283 | 0.277 |

**Supplementary Table 9.** Benchmarking results of species-level taxonomic classification using NGS data (compared with using multiple linear genomes for other tools). In contrast, apart from PanTax, all other tools incorporate multiple linear genomes of each species into the reference database (that is using RefDB:13404), rather than using a single genome as a representative of a specific species (that is using RefDB:8778).

| Method | PanTax | Kraken | Kraken2/Bracken | KrakenUniq | CLARK | Centrifuge | Qmatey | MetaPhlAn4 |
|---|---|---|---|---|---|---|---|---|
| **Index Construction** | | | | | | | | |
| CPU(h) | 17.7 | 32.5 | 162.8 | 35.9 | 10.1 | 22.0 | - | - |
| Wall Time(h) | 5.9 | 4.7 | 3.5 | 6.5 | 10.2 | 2.2 | - | - |
| Memory(G) | 516.8 | 347.7 | 47.2 | 347.7 | 392.2 | 295.6 | - | - |
| **sim-low** | | | | | | | | |
| CPU(h) | 18.3 | 32.7 | 162.8 | 36.1 | 10.3 | 22.9 | 0.4 | 7.3 |
| Wall Time(h) | 6.2 | 6.3 | 3.5 | 8.7 | 10.3 | 2.2 | 0.3 | 0.4 |
| Memory(G) | 490.4 | 297.5 | 49.7 | 298.5 | 128.7 | 16.3 | 9.1 | 18.7 |
| **sim-high** | | | | | | | | |
| CPU(h) | 22.2 | 33.8 | 163.3 | 37.6 | 11.6 | 29.7 | 1.5 | 67.8 |
| Wall Time(h) | 6.3 | 4.9 | 3.6 | 6.8 | 10.7 | 2.3 | 0.9 | 2.3 |
| Memory(G) | 492.4 | 328.8 | 53.9 | 348.5 | 138.9 | 17.7 | 11.2 | 18.7 |
| **CAMI gastrointestinal tract** | | | | | | | | |
| CPU(h) | 19.5 | 33.2 | 163.1 | 36.6 | 11.0 | 23.7 | 0.7 | 3.4 |
| Wall Time(h) | 7.3 | 6.8 | 3.6 | 9.1 | 10.5 | 2.3 | 1.5 | 0.1 |
| Memory(G) | 488.8 | 328.2 | 50.6 | 331.0 | 132.6 | 17.0 | 9.8 | 18.7 |
| **CAMI strain-madness** | | | | | | | | |
| CPU(h) | 19.3 | 32.8 | 162.9 | 36.2 | 10.4 | 25.0 | 0.5 | 2.7 |
| Wall Time(h) | 6.1 | 4.9 | 3.5 | 6.5 | 10.3 | 2.2 | 0.4 | 0.1 |
| Memory(G) | 491.2 | 323.4 | 49.8 | 326.1 | 129.4 | 16.6 | 9.7 | 18.7 |
| **ATCC** | | | | | | | | |
| CPU(h) | 18.8 | 32.6 | 162.9 | 36.1 | 10.3 | 22.8 | 0.3 | 2.2 |
| Wall Time(h) | 6.1 | 6.6 | 3.5 | 7.8 | 10.3 | 2.2 | 0.2 | 0.1 |
| Memory(G) | 491.2 | 183.2 | 49.5 | 231.9 | 128.7 | 16.4 | 8.9 | 18.7 |
| **Zymo1** | | | | | | | | |
| CPU(h) | 19.2 | 32.8 | 162.9 | 36.2 | 10.5 | 24.6 | 0.4 | 1.6 |
| Wall Time(h) | 6.2 | 6.8 | 3.5 | 8.7 | 10.3 | 2.3 | 0.3 | 0.4 |
| Memory(G) | 492.0 | 277.4 | 49.6 | 278.0 | 130.2 | 16.6 | 9.3 | 18.7 |

**Supplementary Table 10.** Runtime and memory usages of benchmarking tools on NGS data. During index construction, we measured the CPU time, wall time, and maximum RAM usage required by the benchmarking tools to build an index utilizing 64 threads. Notably, CLARK was constrained to a single thread for index building. MetaPhlAn2 and MetaPhlAn4 utilized their respective pre-built maker-based databases, while Qmatey did not require index construction. Furthermore, both index construction and querying of Bracken were post-processed based on Kraken2.

| Method | PanTax | Kraken | Kraken2 | KrakenUniq | CLARK | Centrifuge | MetaMaps | MegaBLAST |
|---|---|---|---|---|---|---|---|---|
| **Index Construction** | | | | | | | | |
| CPU(h) | - | 32.5 | 162.8 | 35.9 | 10.1 | 22.0 | 0.1 | 0.5 |
| Wall Time(h) | - | 4.7 | 3.5 | 6.5 | 10.2 | 2.2 | 0.1 | 0.6 |
| Memory(G) | - | 347.7 | 47.2 | 347.7 | 392.2 | 295.6 | 5.4 | 6.1 |
| **sim-low PacBio HiFi** | | | | | | | | |
| CPU(h) | 72.8 | 32.7 | 162.9 | 36.1 | 10.3 | 24.0 | 6.5 | 41.0 |
| Wall Time(h) | 2.9 | 10.8 | 3.6 | 11.1 | 10.3 | 2.7 | 7.0 | 33.5 |
| Memory(G) | 475.9 | 297.1 | 48.8 | 296.1 | 129.6 | 16.3 | 301.6 | 21.3 |
| **sim-low PacBio CLR** | | | | | | | | |
| CPU(h) | 82.5 | 32.7 | 162.9 | 36.1 | 10.3 | 22.6 | 4.0 | 36.6 |
| Wall Time(h) | 3.0 | 10.8 | 3.5 | 14.1 | 10.2 | 2.6 | 7.0 | 33.6 |
| Memory(G) | 474.3 | 328.0 | 48.9 | 328.6 | 129.6 | 16.6 | 300.9 | 21.2 |
| **sim-low ONT R10.4** | | | | | | | | |
| CPU(h) | 71.9 | 32.7 | 162.9 | 36.1 | 10.3 | 24.0 | 6.5 | 39.4 |
| Wall Time(h) | 2.8 | 10.8 | 3.5 | 11.2 | 10.2 | 2.6 | 6.9 | 32.8 |
| Memory(G) | 474.4 | 322.6 | 48.9 | 322.4 | 129.6 | 16.5 | 303.0 | 21.2 |
| **sim-high PacBio HiFi** | | | | | | | | |
| CPU(h) | 395.2 | 34.1 | 163.4 | 37.6 | 11.7 | 25.4 | 81.3 | 408.6 |
| Wall Time(h) | 13.2 | 8.1 | 4.3 | 8.4 | 10.3 | 2.3 | 4.4 | 266.6 |
| Memory(G) | 474.5 | 328.8 | 50.1 | 330.2 | 147.4 | 16.7 | 302.2 | 22.2 |
| **sim-high PacBio CLR** | | | | | | | | |
| CPU(h) | 556.3 | 33.9 | 163.3 | 37.5 | 12.0 | 28.8 | 22.3 | 315.7 |
| Wall Time(h) | 17.4 | 7.8 | 4.0 | 8.3 | 10.3 | 2.3 | 4.7 | 226.1 |
| Memory(G) | 474.7 | 329.1 | 51.8 | 332.2 | 147.4 | 19.4 | 300.9 | 22.3 |
| **sim-high ONT R10.4** | | | | | | | | |
| CPU(h) | 479.3 | 34.0 | 163.3 | 37.7 | 11.4 | 29.8 | 76.6 | 394.9 |
| Wall Time(h) | 14.2 | 7.9 | 4.2 | 8.4 | 10.3 | 2.3 | 6.2 | 259.6 |
| Memory(G) | 475.5 | 329.1 | 50.6 | 330.5 | 147.5 | 18.3 | 303.2 | 22.1 |
| **CAMI gastrointestinal tract** | | | | | | | | |
| CPU(h) | 328.0 | 33.3 | 163.1 | 36.7 | 11.0 | 23.4 | 7.1 | 86.5 |
| Wall Time(h) | 6.8 | 6.7 | 3.6 | 11.9 | 10.5 | 2.2 | 5.5 | 208.1 |
| Memory(G) | 474.6 | 329.0 | 48.7 | 331.6 | 136.6 | 16.6 | 298.0 | 20.4 |
| **CAMI strain-madness** | | | | | | | | |
| CPU(h) | 89.7 | 32.8 | 162.9 | 36.2 | 10.4 | 23.1 | 4.1 | 37.7 |
| Wall Time(h) | 3.8 | 4.7 | 3.5 | 8.7 | 10.3 | 2.3 | 3.7 | 36.7 |
| Memory(G) | 474.9 | 328.0 | 48.2 | 328.4 | 131.0 | 16.3 | 299.5 | 18.3 |
| **ATCC HiFi** | | | | | | | | |
| CPU(h) | 585.1 | 35.6 | 163.8 | 39.1 | 12.4 | 28.5 | 208.3 | - |
| Wall Time(h) | 10.4 | 6.0 | 3.5 | 8.5 | 10.3 | 2.3 | 208.3 | - |
| Memory(G) | 474.8 | 315.7 | 49.4 | 315.7 | 165.8 | 16.8 | 299.7 | - |
| **Zymo1 ONT R9.4.1** | | | | | | | | |
| CPU(h) | 528.6 | 34.3 | 163.6 | 37.8 | 12.7 | 26.6 | 62.5 | 426.7 |
| Wall Time(h) | 9.6 | 6.7 | 3.6 | 7.8 | 10.3 | 2.3 | 8.1 | 362.7 |
| Memory(G) | 475.8 | 329.0 | 48.9 | 329.9 | 154.3 | 19.7 | 300.1 | 36.7 |
| **Zymo1 ONT R10.4** | | | | | | | | |
| CPU(h) | 519.2 | 34.3 | 163.3 | 37.5 | 12.2 | 28.1 | 39.9 | 335.2 |
| Wall Time(h) | 9.4 | 11.0 | 3.5 | 8.6 | 10.3 | 2.3 | 3.2 | 246.1 |
| Memory(G) | 473.9 | 328.9 | 48.9 | 330.1 | 151.3 | 18.6 | 299.8 | 31.6 |
| **Zymo2 HiFi** | | | | | | | | |
| CPU(h) | 498.8 | 34.5 | 163.7 | 38.2 | 12.7 | 27.9 | 107.4 | - |
| Wall Time(h) | 9.3 | 6.7 | 3.5 | 8.2 | 10.4 | 2.3 | 5.9 | - |
| Memory(G) | 474.1 | 286.4 | 48.9 | 288.8 | 160.9 | 16.6 | 299.8 | - |
| **NWC PacBio** | | | | | | | | |
| CPU(h) | 186.5 | 33.2 | 163.0 | 36.6 | 11.0 | 23.7 | 9.9 | 90.2 |
| Wall Time(h) | 4.8 | 6.9 | 3.5 | 8.9 | 10.3 | 2.2 | 5.0 | 72.3 |
| Memory(G) | 476.0 | 322.7 | 48.3 | 323.3 | 136.6 | 17.7 | 298.7 | 19.4 |
| **NWC ONT** | | | | | | | | |
| CPU(h) | 58.2 | 35.2 | 162.9 | 36.1 | 10.3 | 22.5 | 6.6 | 31.4 |
| Wall Time(h) | 2.1 | 7.5 | 3.5 | 8.2 | 10.3 | 2.2 | 3.1 | 23.2 |
| Memory(G) | 475.8 | 280.1 | 48.4 | 282.3 | 129.8 | 17.2 | 298.8 | 16.6 |

**Supplementary Table 11.** Runtime and memory usages of benchmarking tools on TGS data. During index construction, we measured the CPU time, wall time, and maximum RAM usage required by the benchmarking tools to build an index utilizing 64 threads. Notably, CLARK was constrained to a single thread for index building. MegaBLAST was killed after running with 64 threads for more than fifteen days.

### Supplementary Methods

#### Algorithm

**Algorithm 1 graph-based clustering**

**Input:** The genome set $G = \{G_1, G_2, ..., G_N\}$ with N genomes from M species $S = \{S_1, S_2, ..., S_M\}$
**Output:** The non-redundant genome set $R = \{R[S_1], R[S_2], ..., R[S_M]\}$ from M species

```
for S_i ∈ S do
    L[S_i] = ComputeGenomeN50(S_i)
    B[S_i] = GetRepresentaiveGenome(S_i)        // If the species has a reference genome or representative genome in NCBI RefSeq
for S_1 to S_M do
    C_i = SelectGenomes(S_i, number)        // default number 100
    for pair strains x, y in C_i do
        A[x][y] = ComputeANI(x, y)
// main
for S_i ∈ S do
    if S_i has only one strain h then
        R[S_i] = h
    else
        for pair strains x, y ∈ C_i do
            E = [A[x][y] >= 99.9]
        Node = C_i
        Edge = E
        Graph = Graph(Node, Edge)
        H = ConnectedComponents(G)
        for H_j ∈ H do
            for h ∈ H_j do
                if L(h) is max then
                    add h_i to O
        for pair strains x, y ∈ O do
            D = [A[x][y] >= 95]
        Node = O
        Edge = D
        Graph = Graph(Node, Edge)
        K = FindCliques(G)
        K = Sort(K)
        R[S_i] = K_1        // K_1 has the maximum number of strains
        for K_j ∈ K do
            for strain x ∈ K_j do
                if x == B[S_i] then
                R[S_i] = K_j
        if S_i all A[x][y] < 95 then
            if B[S_i] ∈ C_i then
                R[S_i] = B[S_i]
            else
                for h ∈ C_i do
                    if L(h) is max then
                        R[S_i] = h
R[S_i] = SelectGenomes(R[S_i], number)        // default number 10
```

#### AUPR calculation example

We present a straightforward example to elucidate the methodology for calculating AUPR (Area Under the Precision-Recall Curve). Notably, we refrain from utilizing the sklearn Python package for AUPR computation due to its reliance on a Boolean vector that merely indicates the correctness of taxa classification. This approach may lead to biased comparisons as it disregards the ground truth taxa present in the dataset when calculating both precision and recall.

Consider a predicted Boolean vector [1, 0, 1, 0], where '1' signifies a true positive (TP) taxon and '0' denotes a false positive (FP) taxon. The corresponding relative abundance vector, sorted in descending order, is [0.5, 0.3, 0.15, 0.05], and the total number of ground truth strains (TP+FN) is 3.

To calculate precision and recall, we consider taxa above various cutoffs. At a cutoff of 0.05, the updated predicted vector and relative abundance vector are [1, 0, 1, 0] and [0.5, 0.3, 0.15, 0.05], respectively. This yields a precision of 0.5 (2/4) and a recall of 0.67 (2/3), resulting in the first point on the diagram (0.67, 0.5).

Increasing the cutoff to 0.15 updates the vectors to [1, 0, 1] and [0.5, 0.3, 0.15], respectively, leading to a precision and recall of 0.67 (2/3), marking the second point (0.67, 0.67).

At a cutoff of 0.3, the vectors become [1, 0] and [0.5, 0.3], resulting in a precision of 0.5 (1/2) and a recall of 0.33 (1/3), placing the third point at (0.33, 0.5).

Finally, at a cutoff of 0.5, the vectors are [1] and [0.5], yielding a precision of 1 (1/1) and a recall of 0.33 (1/3), marking the fourth point (0.33, 1).

To ensure a smooth diagram, we initiate the plot at (0, 1), where 1 represents the precision of the fourth point. Since the recall of the first point is less than 1, we penalize the AUPR between the greatest recall and 100% recall by setting the endpoint to (0.67, 0), where 0.67 is the recall of the first point. The precision and recall vectors are [1, 1, 0.5, 0.67, 0.5, 0] and [0, 0.33, 0.33, 0.67, 0.67, 1], respectively. We utilize the trapz function from the NumPy Python package to approximate the AUPR by applying the trapezoidal rule.

#### Commands and versions of tools used for comparison

- PanTax v1.0.0
  `bash pantax.sh -f $genomes_info -s -p -r $read.fq --species-level --strain-level` #short read
  `bash pantax.sh -f $genomes_info -l -r $read.fq --species-level --strain-level` #long read

- Kraken v1.1.1

  Index construction
  (1) Download taxonomy files
  `kraken-build --download-taxonomy --db $Kraken_DB --threads 32`
  (2) Prepare all genomes in Kraken2 format and repeat this step to add all genomes to library.
  `kraken-build --add-to-library $genome --db $Kraken_DB`
  (3) Build index with 64 threads , with default parameters.
  `kraken2-build --build --db $Kraken2_DB --threads 64`

  Query
  (1) Short read
  `kraken --db $KrakenDB --output kraken_query_reads --threads 64`
  `--fastq-input --paired $read1.fq $read2.fq`
  (2) Long read
  `kraken --db $KrakenDB --fastq-input $read.fq --output kraken_query_reads --threads 64`
  The kraken_query_reads specifies the assignment of each read, so we can measure read level classification performance and calculate abundance by parsing it. All other parameters are default.

- Kraken2 v2.1.3

  Index construction
  (1) Download taxonomy files
  `kraken2-build --download-taxonomy --db $Kraken2_DB --threads 32`

(2) Prepare all genomes in Kraken2 format and repeat this step to add all genomes to library.
```
kraken2-build --add-to-library $genome --db $Kraken2_DB
```
(3) Build index with 64 threads , with default parameters.
```
kraken2-build --build --db $Kraken2_DB --threads 64
```

Query
(1) Short read
```
kraken2 --db $Kraken2_DB --output kraken2_query_reads --report kraken2_query_report
 --threads 64 --paired $read1.fq $read2.fq
```
(2) Long read
```
kraken2 --db $Kraken2_DB --output kraken2_query_reads --report kraken2_query_report
 --threads 64 $read.fq
```
The kraken2_query_reads specifies the assignment of each read, which can be used to measure the classification performance and calculate the abundance. The kraken2_query_report is used as input for Bracken. All other parameters are default.

- Bracken v2.9
  Index construction
  ```
  bracken-build -d $Kraken2DB -t 64 -l <125/150>
  ```
  Bracken index construction is based on Kraken2 database. It requires read length(-l) for its index construction, so it is not suitable for long read.

  Query
  ```
  bracken -d $KrakenDB -i kraken2_query_report -o bracken_query -r <125/150>
  ```
  Bracken is generally used for short read because it needs to specify the read length. Unlike other tools, Bracken does not take a fq file as input, but post-processes based on the results of kraken2 report file(kraken2_query_report). We calculate the abundance by parsing bracken_query. All other parameters are default.

- KrakenUniq v1.0.4
  Index construction
  (1) Download taxonomy files
  ```
  krakenuniq-download --db $KrakenUniqDB --threads 32 taxonomy
  ```
  (2) Create a subdirectory named "library" in $KrakenUniqDB and copy all genomes to "library". KrakenUniq needs a sequence ID to taxonomy ID mapping for each sequence. We create a mapping file with three tab-separated fields that are, in order, the sequence ID (i. e. the sequence header without '>' up to the first space), the taxonomy ID and the genome or assembly name.
  (3) Build index with 64 threads , with default parameters.
  ```
  krakenuniq-build --db $KrakenUniqDB --jellyfish-bin $jellyfish_path --threads 64
  ```

  Query
  (1) Short read
  ```
  krakenuniq --db $KrakenUniqDB --output krakenuniq_query_reads --report-file
  krakenuniq_query_report --threads 64 --paired $read1.fq $read2.fq
  ```
  (2) Long read
  ```
  krakenuniq --db $KrakenUniqDB --output krakenuniq_query_reads --report-file
  krakenuniq_query_report --threads 64 $read.fq
  ```
  The krakenuniq_query_reads specifies the assignment of each read, which can be used to measure the classification performance and calculate the abundance. All other parameters are default.

- Centrifuge v1.0.4
  Index construction
  ```
  centrifuge-build -p 64 --conversion-table seqid2taxid.map --taxonomy-tree nodes.dmp
  ```

`--name-table names.dmp reference_genomes.fna $centrifugeDB` seqid2taxid.map is a two columns, tab-separated file, which maps each contig to taxonomic ID(mapping sequence IDs to taxonomic IDs as KrakenUniq's mapping file). The nodes.dmp and names.dmp was obtained from Kraken2 first step. reference_genomes.fna is a fasta file that concatenates all input genomes.

Query
(1) Short read
```
centrifuge -k 1 -x $centrifugeDB -1 $read1.fq -2 $read2.fq -S centrifuge_query_reads
--report-file centrifuge_query_report --threads 64
```
(2) Long read
```
centrifuge -k 1 -x $centrifugeDB -U $read.fq -S centrifuge_query_reads
--report-file centrifuge_query_report --threads 64
```
The centrifuge_query_reads specifies the assignment of each read, which can be used to measure the classification performance and calculate the abundance. All other parameters are default.

- CLARK v1.2.6.1
  Index construction and query
  We use CLARK executable, which can directly complete index construction and query together, unlike other tools. If the database clarkDB does not exist at the first run, the index will be built and the query will be executed. Otherwise, the query will be executed directly.
  (1) short read
  ```
  CLARK -T genome_map_clark.txt -D $clarkDB -P $read1.fq $read2.fq -R CLARK_query_reads -n 64
  ```
  (2) long read
  ```
  CLARK -T genome_map_clark.txt -D $clarkDB -O $read.fq -R CLARK_query_reads -n 64
  ```
  genome_map_clark.txt is a two columns, tab-separated file, which maps each genome(fasta file) to taxonomic ID. The CLARK_query_reads specifies the assignment of each read, which can be used to measure the classification performance and calculate the abundance. All other parameters are default.

- MegaBLAST v2.12.0
  Index construction
  ```
  makeblastdb -in reference_genomes.fna -dbtype nucl -parse_seqids -taxid_map readsid_taxid.txt
  -out $megablastDB
  makembindex -input $megablastDB -iformat blastdb
  ```
  reference_genomes.fna is a fasta file that concatenates all input genomes as Centrifuge. readsid_taxid.txt is a two columns, tab-separated file, which maps each contig(represented by taxonomic ID) to species taxid.
  Query
  ```
  blastn -db $megablastDB -query $read.fq -out megablast_result -max_hsps 10 -num_threads 64
  -task megablast -use_index true -outfmt "6 qseqid sseqid pident length mismatch qstart qend
  sstart send evalue bitscore staxids"
  ```
  The megablast_result specifies the assignment of each read, which can be used to measure the classification performance and calculate the abundance. But a read has multiple matching results in megablast_result, we only select the first one. All other parameters are default.

- MetaMaps v0.1
  Index construction
  ```
  perl combineAndAnnotateReferences.pl --inputFileList taxid2seq.txt
  --outputFile reference_genomes.fa --taxonomyInDirectory taxonomy
  --taxonomyOutDirectory taxonomy_out
  perl buildDB.pl --DB $metamapsDB --FASTAs reference_genomes.fa --taxonomy taxonomy_out
  ```
  taxid2seq.txt is a two columns, tab-separated file, which maps taxonomic ID to each genome(fasta file). The taxonomy information downloaded from NCBI.
  Query
  ```
  metamaps mapDirectly -r reference_genomes.fa -q $read.fq -t 64 -o classification_results
  ```

```
metamaps classify --mappings classification_results --DB $metamapsDB -t 64
```
The classification_results.EM.reads2Taxon specifies the assignment of each read, which can be used to measure the classification performance and calculate the abundance. All other parameters are default.

- MetaPhlAn2 v2.6.0
  Index construction
  ```
  bowtie2-build mpa_v20_m200.fna mpa_v20_m200
  ```
  mpa_v20_m200.fna was downloaded from MetaPhlAn database.
  Query
  ```
  python metaphlan2.py $read1.fq,$read2.fq --input_type fastq -o abundance.txt --nproc 64
  ```
  Unlike other tools, MetaPhlAn2 directly outputs abundance. Note that MetaPhlAn2 can only be used for short read. All other parameters are default.

- MetaPhlAn4 v4.0.6
  Query
  ```
  metaphlan $read1.fq,$read2.fq --nproc 64 --input_type fastq -o abundance.txt
  ```
  MetaPhlAn4 also directly outputs abundance as MetaPhlAn2. Note that MetaPhlAn4 can only be used for short read. All other parameters are default.

- Qmatey v0.5.1
  Index construction and query
  Qmatey runs index construction and query together. It works in the specified directory, which provides the necessary files listed below. Qmatey running needs to prepare input_dbfasta and map_taxids file, and provides their file path for config.sh. input_dbfasta is a fasta file that concatenates all input genomes. map_taxids is a two columns, tab-separated file, which maps each contig to taxonomic ID. Paired-end reads need to be prepared in the subdirectory samples/pe under this directory.
  ```
  sh Qmatey $directory
  ```
  All other parameters are default. The result in *_taxainfo_mean_normalized.txt, and abundance calculated by normalization. Note that the results of Qmatey do not seem to be stable.

- StrainEst
  Index construction
  The StrainEst does not provide a script for automating the construction of the database, so we implement it step by step according to the description of its paper.
  ```
  strainest mapgenomes A1.fasta A2.fasta... SR.fasta MA.fasta
  strainest map2snp SR.fasta MA.fasta snp.dgrp
  strainest snpdist snp.dgrp snp_dist.txt hist.pdf
  strainest snpclust snp.dgrp snp_dist.txt snp_clust.dgrp clusters.txt
  bowtie2-build MA.fasta MA
  ```
  The specific description of commands and parameters is in the paper, which is only briefly described here. SR.fasta is the reference genome or representative genome of specified species in NCBI RefSeq. Here we select all the genomes as the representative genomes of the tool, rather than only some for metagenome alignment.
  Query
  ```
  sickle pe -f $read1.fq -r $read2.fq -t sanger -o reads1.trim.fastq -p reads2.trim.fastq
  -s reads.singles.fastq -q 20
  bowtie2 --very-fast --no-unal -x MA -1 reads1.trim.fastq -2 reads2.trim.fastq -S reads.sam
  samtools view -b reads.sam > reads.bam
  samtools sort reads.bam -o reads.sorted.bam
  samtools index reads.sorted.bam
  strainest est snp_clust.dgrp reads.sorted.bam $outputdir
  ```
  All other parameters are default. The abundance output of all genomes is in abund.txt.

- StrainGE
  Index construction
  For every genome, firstly run `straingst kmerize -o $strainge_db/genome.hdf5 $genome`
  `straingst kmersim --all-vs-all -t 64 -S jaccard -S subset $strainge_db/*.hdf5 > similarities.tsv`
  `straingst cluster -i similarities.tsv -d -C 0.99 -c 0.90 --clusters-out clusters.tsv $strainge_db/*.hdf5 > references_to_keep.txt`
  `straingst createdb -f references_to_keep.txt -o pan-genome-db.hdf5`
  All genomes are prepared in specified directory as strainge_db.
  Query
  `straingst kmerize -k 23 -o result.hdf5 $read1.fq $read2.fq`
  `straingst run -O -o result pan-genome-db.hdf5 result.hdf5`
  All other parameters are default. We only run the first step of StrainGE, StrainGST, to obtain the abundance. The abundance output of possible genomes is in result.strains.tsv.

- StrainScan
  Index construction
  `strainscan_build -i $input_fasta_dir -o $strainscan_db -t 64`
  All genomes are prepared in specified directory as $input_fasta_dir
  Query
  `strainscan -i $read1.fq -j $read2.fq -d $strainscan_db -o strainscan_result`
  All other parameters are default. The abundance output of possible genomes is in final_report.txt. Possible genomes abundance is obtained by normalized the `Predicted_Depth (Ab*cls_depth) column`.
